# Positive Selection and Heat-Response Transcriptomes Reveal Adaptive Features of the Brassicaceae Desert Model, *Anastatica hierochuntica*

**DOI:** 10.1101/2021.05.23.445339

**Authors:** Gil Eshel, Nick Duppen, Guannan Wang, Dong-Ha Oh, Yana Kazachkova, Pawel Herzyk, Anna Amtmann, Michal Gordon, Vered Chalifa-Caspi, Michelle Arland Oscar, Shirli Bar-David, Amy Marshall-Colon, Maheshi Dassanayake, Simon Barak

**Author notes:** Authors for correspondence: Simon Barak.; Maheshi Dassanayake. These authors contributed equally to the article (Gil Eshel, Nick Duppen). Department of Plant and Environmental Sciences, Weizmann Institute of Science, Rehovot 7610001, Israel (Yana Kazachkova).

## Abstract

- Plant adaptation to a desert environment and its endemic heat stress is poorly understood at the molecular level. The naturally heat-tolerant Brassicaceae species *Anastatica hierochuntica* is an ideal extremophyte model to identify genetic adaptations that have evolved to allow plants to tolerate heat stress and thrive in deserts.
- We generated an *A. hierochuntica* reference transcriptome and pinpointed extremophyte adaptations by comparing *Arabidopsis thaliana* and *A. hierochuntica* transcriptome responses to heat and identifying positively selected genes in *A. hierochuntica*.
- The two species exhibit similar transcriptome adjustment in response to heat and the *A. hierochuntica* transcriptome does not exist in a constitutive heat “stress-ready” state. Furthermore, the *A. hierochuntica* global transcriptome as well as heat-responsive orthologs, display a lower basal and higher heat-induced expression than in *A. thaliana*. Genes positively selected in multiple extremophytes are associated with stomatal opening, nutrient acquisition, and UV-B induced DNA repair while those unique to *A. hierochuntica* are consistent with its photoperiod-insensitive, early-flowering phenotype.
- We suggest that evolution of a flexible transcriptome confers the ability to quickly react to extreme diurnal temperature fluctuations characteristic of a desert environment while positive selection of genes involved in stress tolerance and early flowering could facilitate an opportunistic desert lifestyle.

## Introduction

Plant species inhabiting extreme environments - so-called extremophytes - are able to thrive in some of the most inhospitable environments on Earth that are characterized by severe abiotic stresses. These stresses include drought and temperature extremes in deserts, intense cold in the Antarctic, and habitats both on land and in the sea that are typified by acute salinity (Amtmann, 2009; John et al., 2009; Dassanayake et al., 2010; Oh et al., 2012; Lawson et al., 2014; Farrant et al., 2015; Fan et al., 2018; Kazachkova et al., 2018; Oscar et al., 2018). Understanding the molecular mechanisms by which extremophytes adapt to their stressful environments could aid in identifying targets for molecular breeding efforts to improve crop stress tolerance, as well as facilitate the development of extremophyte- based agriculture (Bressan et al., 2013; Shabala, 2013; Cheeseman et al., 2015; Ventura et al., 2015). While tolerance to salt stress has been extensively investigated in halophytes that are adapted to highly saline environments (Flowers et al., 2015; Kazachkova et al., 2018; Wang et al., 2021a), our understanding of plant molecular adaptations to stresses characteristic of desert habitats is still in its infancy (e.g. Granot et al., 2009; Yates et al., 2014; Oh et al., 2015; Obaid et al., 2016; Eshel et al., 2021; Wan et al., 2021). Yet, desert species could represent a treasure trove of molecular determinants that confer tolerance to the multiple stresses such as drought, salinity, low soil nutrient levels, and heat stress. With global temperatures projected to continue rising (IPCC, 2021), plant adaptation to heat stress is of particular importance due to its negative effects on plant physiology, particularly at the reproductive stage, leading to severe consequences upon yield (Mittler and Blumwald, 2010; Lesk et al., 2016; Chaturvedi et al., 2021; Wang et al., 2021b). Indeed, tolerance to heat stress in wheat at the reproductive stage has been identified as a key trait to increase yield potential under projected climate change (Stratonovitch and Semenov, 2015).

To gain insight into genetic adaptations that facilitate an extremophyte lifestyle, comparative physiological and molecular analyses of stress-sensitive *Arabidopsis thaliana* and its Brassicaceae extremophyte relatives have proven to be a powerful approach (Kraemer, 2010; Koenig and Weigel, 2015; Kazachkova et al., 2018). Indeed, these extremophyte relatives are becoming premier models for understanding plant adaptation to extreme environments with the development of a number of genetic resources including chromosome-level genome assemblies, natural accession collections, transformation protocols, and web resources (http://extremeplants.org/) (Zhu et al., 2015; Kazachkova et al., 2018; Wang et al., 2019). Yet, an extremophyte Brassicaceae model that represents desert species has not hitherto been developed. Such a model could leverage the functional genomics knowledge that exists for *A. thaliana* thereby facilitating comparative analyses to understand plant adaptations to the extreme desert environment. We have therefore been studying the *A. thaliana* relative, *Anastatica hierochuntica* L., also known as the ‘True Rose of Jericho’, a Saharo-Arabian desert species (Fig. 1A) which also occupies the uppermost, driest zones of wadies or runnels of the Israeli Negev desert (Friedman and Stein, 1980; Friedman et al., 1981; Fig. 1B). This arid region has temperatures varying between -3.6 and 46 °C, an annual rainfall between 25 and 200 mm, and soil nitrate levels ranging from 0.4 to 4 mM (Gutterman, 2002; Ward, 2009; Eshel et al., 2017). We have demonstrated that *A. hierochuntica* is highly tolerant to heat, low soil nitrogen, and moderately tolerant to salt stress (Eshel et al., 2017). Moreover, the plant exhibits salt-resilient photochemistry and displays constitutively higher levels of metabolites that have a role in scavenging reactive oxygen species, than *A. thaliana* (Eppel et al., 2014; Eshel et al., 2017).

**Figure 1.**
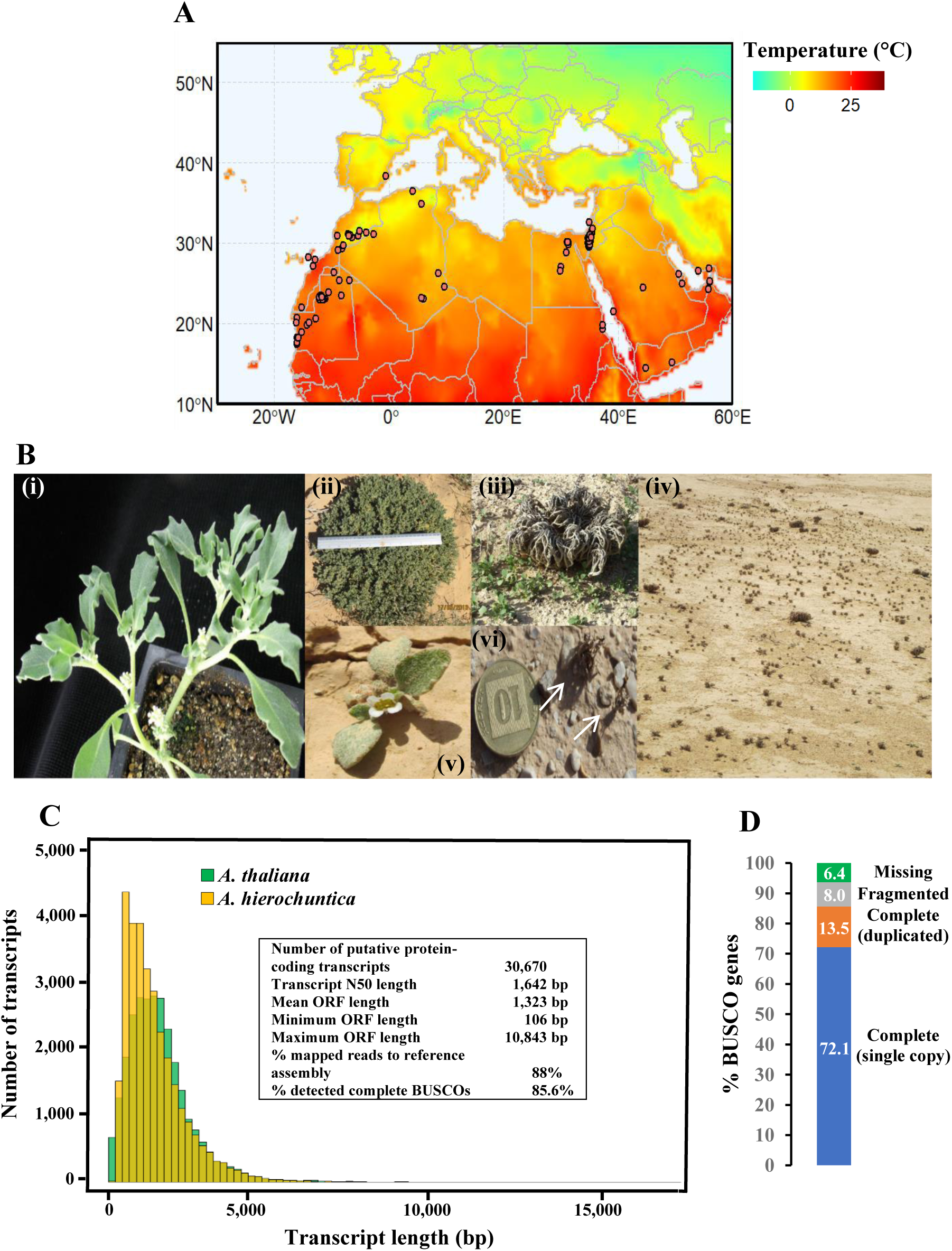
Geographic distribution of *A. hierochuntica* and *de-novo* reference transcriptome. (A), Geographic distribution data are based on *Anastatica* L. in the Global Biodiversity Information Facility database (GBIF Secretariat (2019). GBIF Backbone Taxonomy. Checklist dataset https://doi.org/10.15468/39omei accessed via GBIF.org on 2021-03-12). Average temperature data for this region are from 1948 to Feb 2021 acquired by the Physical Sciences Laboratory (Fan and van den Dool, 2008); (B), Lab-grown and wild *A. hierochuntica* plants. Panels: (i) 40 d-old lab-grown plant. Note the axillary inflorescence at each branch point; (ii) large mature plant from the Ovda valley population in the Negev desert. Ruler length = 30 cm; (iii) young seedlings growing near the dead mother plant from a Neot Smadar population in the Negev desert; (iv) a large population of *A. hierochuntica* in the Ovda valley with high variation in plant size due to spatial and temporal variations in water availability; (v) *A. hierochuntica* seedling already beginning to flower after producing four true leaves (Neot Smadar); (vi) two tiny dead plants (white arrows) from a population near the Dead Sea valley, having already dispersed their few seeds; (C), Transcript length distribution and *A. hierochuntica* assembly descriptive statistics; (D), Assessment of reference transcriptome assembly completeness using the Benchmarking Universal Single-Copy Orthologs (BUSCO) tool (Simão et al., 2015). The percentages of 1,375 single-copy genes, conserved among land plants, identified in the *A. hierochuntica* transcriptome are shown.

In the current study, we assembled an *A. hierochuntica* reference transcriptome and used this resource for two approaches to identify adaptations to an arid environment in a desert annual extremophyte. In the first approach, comparative analysis of heat-response transcriptomes revealed an *A. hierochuntica* transcriptome that is more reactive to heat than that of *A. thaliana*. In the second approach, positive selection analysis identified genes that could contribute to adaptation to extreme conditions in general, and those that could facilitate an opportunistic desert lifestyle.

## Materials and Methods

For all analyses, detailed methods are provided in Supporting Information Methods S1.

### Plant material and growth conditions

Plants for *de novo* reference transcriptome sequencing were grown on MS (Murashige and Skoog, 1962) plates for 5 d in the growth room (16 h light [150 µmol photons m^-2^ s^-1^]/8 h dark; 22 °C). For Illumina sequencing, plate-grown seedlings were directly used. For Roche 454 sequencing, plate-grown seedlings were transferred to pots containing *A. thaliana* soil growth medium (Weizmann Institute of Science), and kept in the growth room until plants developed four true fully-expanded leaves. These plants were then treated as follows: (a) Control (field-capacity, 22 °C); (b) Drought stress (25% field capacity for 1 week); (c) Salt shock (200 mM NaCl in the fertilizer solution), harvested after 1, 3 and 6 h; (d) Heat shock (45 °C), harvested after 0.5, 1, and 2 h. Roots, shoots and flowers (where available) from these soil-grown plants, were harvested separately and snap-frozen in liquid nitrogen. In addition, mature seeds, were imbibed in H_2_O for 8.5 h.

For RNA-seq heat stress experiments, *A. thaliana* and *A. hierochuntica* were grown on plates until cotyledons were fully expanded before transfer to pots containing *Arabidopsis* nitrogen-less soil (Weizmann Institute of Science) and irrigation to field capacity with a custom-made fertilizer solution. After 6 d in the growth room, uniform plants were transferred to two growth chambers (KBWF 720, BINDER GmbH, Tuttlingen, Germany) (16 h light/8 h dark; 23 °C; 60% relative humidity; sunrise, 0.5 h at 100 μmol photons m^-2^ s^-1^; daytime, 250 μmol m^-2^ s^-1^; sunset, 0.5 h at 150 μmol photons m^-2^ s^-1^). Ten days after transfer to soil, heat treatment was initiated in one chamber comprising 3 d at 40/25 °C, day/night temperatures, followed by 2 d recovery at control conditions (Fig. 2A). The other chamber was kept as the control (23 °C). For each condition, three biological replicates comprising 6 pooled plants per replicate (54 samples in total) were used for downstream analyses.

**Figure 2.**
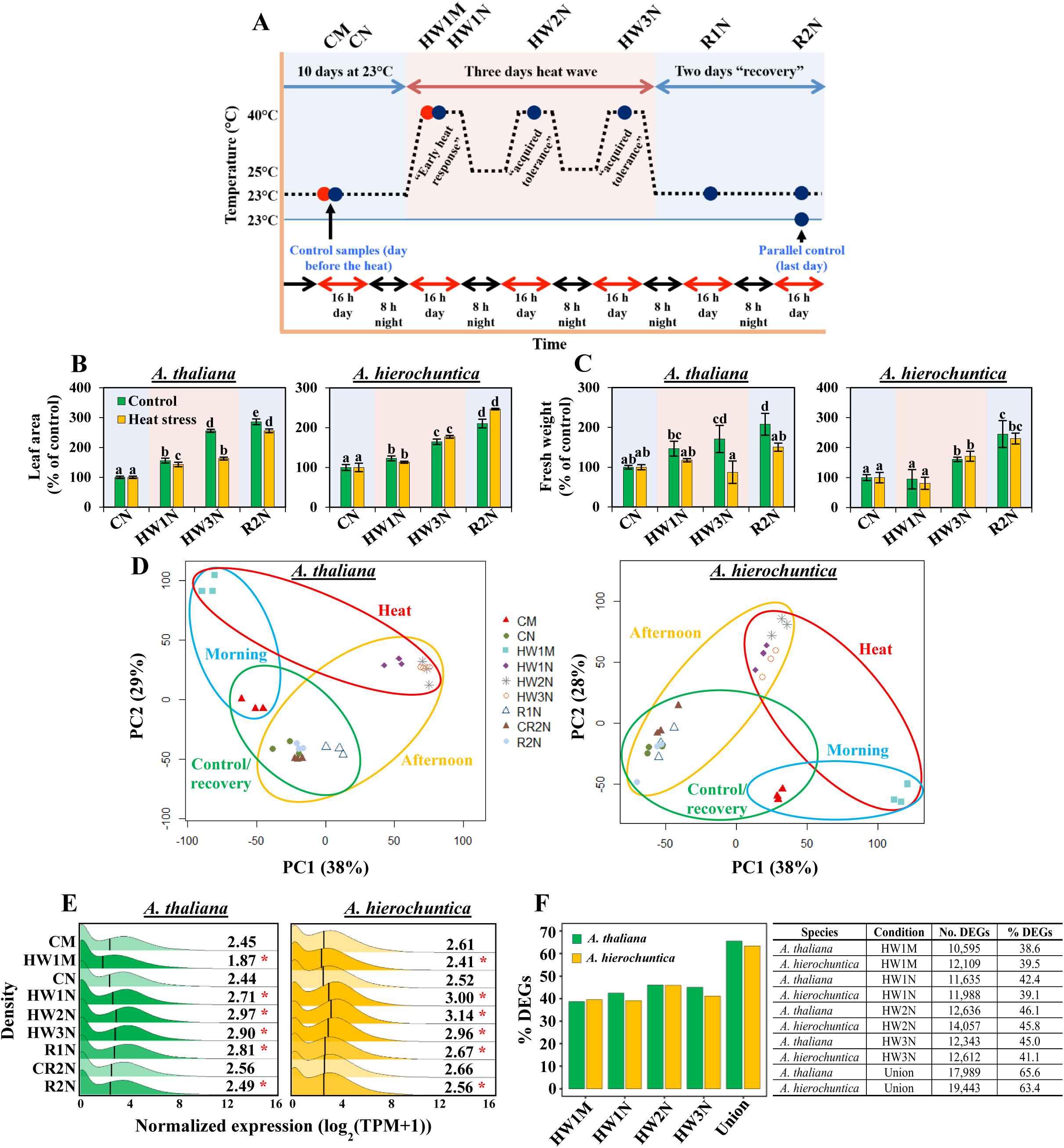
*A. thaliana* and *A. hierochuntica* exhibit similar transcriptome adjustment to heat stress. (A), Experimental design for *A. thaliana* and *A. hierochuntica* control and heat stress conditions. Control plants were harvested the day before the initiation of heat stress and on the last day of the experiment (indicated by vertical arrows) from a parallel 23 °C control chamber. Red and blue circles represent samples harvested 1.5 h (morning) or 7 h (afternoon) respectively, after onset of light/heat. Each circle represents 3 independent experiments, each comprising 6 pooled plants. (B and C), Effect of heat stress on *A. thaliana* and *A. hierochuntica* leaf area (B) and fresh weight (C). Data are mean ± S.D. (n = 5) and are representative of two independent experiments. Letters above bars indicate significant difference at *p* < 0.05 (Tukey HSD test). Blue shading, control conditions; Pink shading, heat conditions. (D), Principal component analysis (PCA) of *A. thaliana* and *A. hierochuntica* transcript levels. Each point represents one biological replicate and the three replicates for each condition are depicted with the same symbol. Symbols are explained in the legend box and refer to the experimental design shown in (A); (E), Comparison of the abundance of 27,416 and 30,670 protein-coding *A. thaliana* and *A. hierochuntica* transcripts, respectively. Asterisks represent significant difference at *p* < 0.05 (Wilcoxon rank sum test) between the treatment compared to its respective control. Black vertical line within plots is median expression; (F), Percent of *A. thaliana* and *A. hierochuntica* differentially expressed genes (DEGs) in response to heat stress. In total, 17,989 *A. thaliana* and 19,443 *A. hierochuntica* genes were differentially expressed in response to heat stress in at least one condition (Dataset S2), and % DEGs was calculated based on 27,416 and 30,670 protein-coding genes for *A. thaliana* and *A. hierochuntica*, respectively. CM, control morning; CN, control afternoon; HW1M, heat wave 1 morning; HW1N, heat wave 1 afternoon; HW2N, heat wave 2 afternoon; HW3N, heat wave 3 afternoon; R1N, day 1 recovery from heat stress afternoon; CR2N, control plants parallel to the R2N time point afternoon; R2N, day 2 recovery from heat stress afternoon; Union, DEGs identified under either HW1M or HW1N or HW2N or HW3N.

### Reference transcriptome and RNA-seq

RNA libraries were prepared using samples suitable for either the *A. hierochuntica* reference transcriptome or the RNA-seq transcriptome analysis:

(i) For the reference transcriptome, equal amounts of total RNA from all samples (i.e. control, various stresses, tissues, time points, see “Plant material and growth conditions”) were pooled and sent to the GenePool genomics facility at the University of Edinburgh, UK for 454 sequencing of a normalized cDNA library. In addition, total RNA from plate-grown seedlings was sent to the Glasgow Polyomics Facility at the University of Glasgow, UK for Illumina sequencing. The reference transcriptome was assembled using a hybrid assembly approach that utilized both Illumina and 454 reads and was annotated based on public databases (Methods S1).
(ii) For RNA-seq, total RNA was extracted from control and heat-treated samples (Fig. 2A) and delivered to the Roy J. Carver Biotechnology Center, University of Illinois, Urbana-Champaign, USA. Libraries were prepared with the Illumina TruSeq Stranded mRNA Sample Prep Kit (Illumina) and 100 bp HiSeq2500 Illumina single-end reads were uniquely mapped to *A. thaliana* TAIR 10 or the *A. hierochuntica* reference transcriptome using the Trinity align_and_estimate_abundance.pl script, which applies the RSEM program (Grabherr et al., 2011; Li and Dewey, 2011) with the Bowtie aligner.

Differentially expressed genes (DEGs) were identified using DESeq2 (Love et al., 2014; Methods S1). For raw read counts and DEGs identified in each species and for various functional groups, see Dataset S2..

Ortholog pairs (17,962; Methods S1) were assigned to the five idealized modes of expression in response to heat stress (Fig. 3A), using Weighted Gene Co-expression Network Analysis (WGCNA; Langfelder and Horvath, 2008) to cluster normalized and quantified expression data into modules containing genes with similar expression profiles (Fig. S3; Dataset S3; Methods S1). For direct comparison of absolute orthologous transcript levels, TPM values of minimum or maximum expression were analyzed for statistically significant difference (*p* ≤ 0.05) with a Student *t*-test.

**Figure 3.**
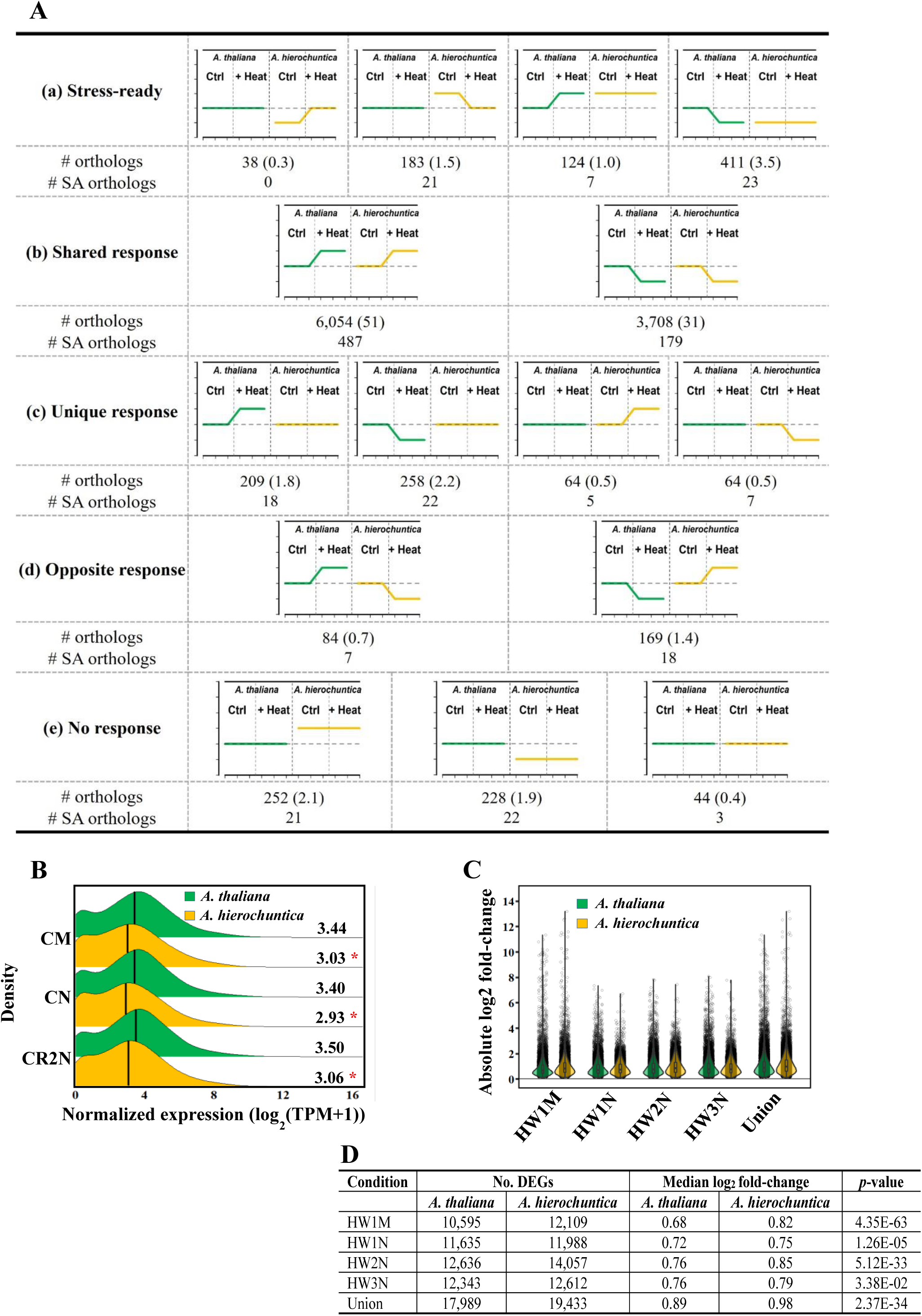
The *A. hierochuntica* transcriptome does not exist in a “stress-ready” state but exhibits a lower basal expression and higher fold-change expression than *A. thaliana* in response to heat stress. (A), Modes of expression of ortholog pairs between *A. thaliana* and *A. hierochuntica* in response to heat stress. WGCNA followed by DESeq2 was used to assign orthologs to response modes (Fig. S3; Dataset S3). Differences in absolute transcripts levels were identified by comparing TPM minimum or maximum expression values (Student’s *t*-test, *p* ≤ 0.05). The green (*A. thaliana*) and orange (*A. hierochuntica*) lines indicate idealized expression patterns of the ortholog pairs in each species under control and heat conditions, compared to the *A. thaliana* control (dashed line). Ctrl, control; + heat, heat stress treatment; SA, genes associated with GO-terms for abiotic stress (Methods S1). Numbers in parentheses, percent of ortholog pairs relative to the total number of orthologs (11,890) assigned to a response mode; (B), Transcript abundance of 17,962 *A. thaliana* and *A. hierochuntica* ortholog pairs under control conditions. (C), Combined violin and box plots showing absolute log_2_ fold-changes of *A. thaliana* and *A. hierochuntica* DEGs in response to heat stress (Dataset S2). The median log_2_ fold-change is shown as a black square inside each box plot; (D), Number of DEGs, median log_2_ fold-change values and *p*-values for panel C. CM, control morning; CN, control afternoon; CR2N, control plants parallel to the R2N (day 2 recovery from heat stress afternoon) time point; HW1M, heat wave 1 morning; HW1N, heat wave 1 afternoon; HW2N, heat wave 2 afternoon; HW3N, heat wave 3 afternoon; Union, DEGs identified under either HW1M or HW1N or HW2N or HW3N. Asterisks represent significant difference at *p* < 0.05 (Wilcoxon rank sum test) between *A. thaliana* and *A. hierochuntica*.

### Phylogenomics and positive selection analysis

To identify orthologous genes among species and generate a maximum likelihood phylogenomic tree, we used coding sequences of the *A. hierochuntica* reference transcriptome and 16 sequenced Brassicaceae species (Dataset S12) with the Agalma phylogenomics pipeline (Dunn et al., 2013).

To detect positive selection, we used the Branch-Site model in the PAML v4.8, CODEML program (Yang, 1997; Yang, 2007). Ortholog groups with sequence representation in at least two of the five extremophytes, were selected to ensure sufficient statistical power (Anisimova et al., 2001). The tested branch(s) were labeled (foreground), and the log likelihood of two models (M1a and M2a), were calculated for each ortholog group. A Likelihood Ratio Test was performed (with Χ^2^ distribution), to identify genes with log likelihood values significantly different between the two models, indicative of deviation from neutral selection. Ortholog groups with a portion of sites in the foreground branches, that had an estimated dN/dS ratio greater than 1, were considered under positive selection. To account for multiplicity, a Benjamini–Yekutieli false discovery rate (FDR) correction (Benjamini and Yekutieli, 2001) was applied using the “qvalue” R package, with a *q*-value < 0.05 cutoff for a gene to be considered as positively selected. Sites under positive selection were identified using the empirical Bayes approach with a posterior probability *p* > 0.95.

For each analysis, different branches on the tree were tested (labeled as foreground) compared with all other branches (background): (i) labeling the external branches of all five extremophyte species as the foreground (4,723 ortholog groups); (ii) labeling the *A. hierochuntica* external branch as the foreground (3,093 ortholog groups); (iii) labeling the *E. salsugineum* external branch as the foreground (4,457 ortholog groups); (iv) labeling the *S. parvula* external branch as the foreground (4,369 ortholog groups); and (v) labeling the *A. thaliana* external branch as the foreground (5,513 ortholog groups). *A. thaliana* was considered as a control/comparator species sensitive to abiotic stresses (Kazachkova et al., 2018). The Venn diagram comparing positive selected genes (Fig. 6B) was generated using an online tool: http://bioinformatics.psb.ugent.be/webtools/Venn/).

Significant positively selected genes as well as all DEGs were tested for enriched GO terms (Fisher’s exact test, with a *q*-value < 0.05 cutoff) using AgriGO (Du et al., 2010), where the *A. thaliana* genome served as background. GOMCL (Wang et al., 2020) was used to summarize non-redundant functional groups).

## Results

### *De novo* assembly and annotation of the *Anastatica hierochuntica* reference transcriptome

To generate a high-quality *A. hierochuntica* reference transcriptome that maximizes coverage of genes, we sequenced and assembled transcripts using RNA pooled from multiple plant organs (root, shoot, flower, seeds), at different developmental stages (early seedling stage, and mature plants before and after flower initiation), and under control, heat, drought and salinity stress conditions (Fig. S1; Methods S1). We identified 30,670 putative protein-coding genes out of the high-confidence 36,871 assembled transcripts (Fig. 1C; Methods S1), and the distribution of transcript lengths was similar to that of *A. thaliana* cDNAs (Fig. 1C). High completeness of the reference transcriptome was evidenced by the detection of 93.6% BUSCOs (Fig. 1D; Simão et al., 2015), comparable to other *de novo* assembled Brassicaceae transcriptomes (Lopez et al., 2017). Furthermore, we obtained 88% sequenced reads that mapped back to the assembled reference transcriptome. These data indicate a high-quality reference transcriptome appropriate for downstream analyses. We annotated the reference transcriptome using sequence similarity to protein databases including NCBI, InterPro, and KEGG (Dataset S1) resulting in annotation of 96% of our assembled transcripts to a known sequence in the reference databases (Methods S1).

### *A. thaliana* and *A. hierochuntica* global transcriptomes exhibit similar adjustment in response to heat stress

Analysis of stress-induced transcriptome responses of halophytic Brassicaceae models suggests that they exist in a “stress-ready” state (Kazachkova et al., 2018; Wang et al., 2021a). However, it is unknown whether a “stress-ready” transcriptome state is the default for all extremophytes or whether plants evolving under different extreme environments exhibit alternate modes of adaptation. Therefore, to test whether a desert species exists in a “stress-ready state”, we performed a comparative analysis of the *A. thaliana* and *A. hierochuntica* transcriptome response to heat stress in young plants at similar developmental stages. Israel Meteorological Service temperature data near *A. hierochuntica* populations during their growing season showed diurnal minimum/maximum night/day temperatures of ∼25 °C and ∼40 °C, respectively (Fig. S2). Thus, to simulate an ecologically relevant scenario with heat treatments that *A. thaliana* plants could also survive (Hayes et al., 2021), plants were exposed to similar three consecutive daily heat waves covering the early heat response and acquired heat tolerance phases (Lindquist, 1986; Hong and Vierling, 2000), with day/night temperatures of 40 °C/25 °C followed by 2 days recovery at 23 °C (Fig. 2A). To minimize shocks, temperatures were gradually ramped up and down at sunrise and sunset, respectively (Methods S1). Control plants were maintained at 23 °C. Plants were harvested either in the morning (1.5 hours after the onset of the light/heat period, red circles in Fig. 2A) or in the afternoon (7 hours after the onset of the light/heat period, blues circles in Fig. 2A). Plants were well-watered throughout the entire experiment to avoid any dehydration effects that could arise due to the heat treatment.

Heat stress had no significant effect on *A. hierochuntica* leaf area in contrast to *A. thaliana* where growth in leaf area was significantly retarded by heat stress although it had almost recovered to control levels 2 days after the end of the heat treatment (Fig. 2B). *A. thaliana* shoot fresh weight was also significantly reduced by heat stress but did not recuperate after 2 days recovery under control conditions while *A. hierochuntica* fresh weight was not affected by heat stress (Fig. 2C). These results illustrate that *A. hierochuntica* is highly tolerant to heat stress and confirm our previous *in vitro* experiments (Eshel et al., 2017).

Transcriptomes of both species under elevated temperature were clearly distinct from those in control conditions (Fig. 2D). The control and heat-stressed samples harvested in the morning were positioned separately from the samples harvested in the afternoon, which could be due to differences in early vs. late heat-mediated gene expression or/and diurnal changes in gene expression. Transcriptomes of plants recovering from heat stress clustered near control noon samples suggesting that, overall, the transcriptomes return to pre-stress conditions. Because both species underwent transcriptional adjustment in response to heat stress, we examined the median expression level across the whole transcriptome for each condition. Compared to their respective controls (CM, CN, CR2N), the median transcript abundance (and total abundance as depicted by the distribution) of both species decreased under heat stress in the morning samples, increased in response to heat treatments in the noon samples, and decreased during recovery (Fig. 2E). Furthermore, the percentage of differentially expressed genes (out of the total number of protein-coding genes) was similar for both species under all heat conditions (Fig. 2F; Dataset S2). These data show that the *A. thaliana* and *A. hierochuntica* global transcriptomes adjust to heat stress with a similar magnitude.

### The *A. hierochuntica* heat-response transcriptome does not exist in a “stress-ready” state

To test our contention that *A. hierochuntica* transcriptome is not “stress-ready”, we used Weighted Gene Co-expression Network Analysis (WGCNA) to identify five types of idealized transcriptional response modes among the expression patterns of 17,962 orthologous pairs from each species (Wang et al., 2021a; Fig. S3; Dataset S3): (a) “Stress-ready” where transcript level under control conditions in one species is equal to the ortholog transcript level under heat in the other species; (b) “Shared response” where expression of both orthologs exhibit a similar response to heat (i.e. both upregulated or downregulated by heat); (c) “Unique response” where expression of an ortholog exhibits a heat response specifically in one species but not in the other; (d) “Opposite response” where expression of the ortholog in one species shows the opposite response in the other species; (e) “No response” where expression of both orthologs does not respond to heat. Of the orthologs categorized within the five response modes, only 4.4% of the orthologs belonged to the “No response” mode (Fig. 3A). The vast majority (82%) of orthologs displayed a shared response mode while about 5% exhibited a unique response and 2.1% showed an opposite response. Importantly, while 535 (4.5%) genes did exhibit a “stress-ready” mode in *A. hierochuntica*, we also detected 221 (1.9%) *A. thaliana* genes displaying a “stress-ready” mode. Similar results were obtained when we examined only orthologs associated with GO terms for abiotic stress responses (Fig. 3A; Methods S1). Taken together, our data show that: (i) the *A. thaliana* and *A. hierochuntica* global transcriptomes adjust to heat stress with a similar magnitude (ii) only a low proportion of genes exhibit a “stress-ready” mode of expression in both species. Thus, our findings do not support a globally “stress-ready” *A. hierochuntica* transcriptome.

### *A. hierochuntica* heat-regulated genes display a higher fold-change and/or lower basal expression compared to *A. thaliana*

Under control conditions, we observed that median basal expression of the *A. hierochuntica* transcriptome was significantly lower than in *A. thaliana* (Fig. 3B). Moreover, differentially expressed genes (DEGs) from the extremophyte displayed a greater heat-mediated fold-change in expression than *A. thaliana* DEGs (Figs. 3C, 3D) suggesting a more reactive heat-response transcriptome. To support these findings, we compared orthologous expression of specific functional groups that exhibited either a shared or unique response mode to heat stress (Dataset S2). Orthologs associated with GO-terms for abiotic stress whose expression displayed shared upregulation by heat exhibited an average lower basal expression in *A. hierochuntica* compared to *A. thaliana* and no significant difference in average % induction of expression (Fig. 4A). Abiotic stress-associated orthologs showing shared heat-mediated downregulated expression displayed both a lower basal and higher % reduction in expression in *A. hierochuntica* compared to *A. thaliana* (Fig. 4B). Similarly, both heat-mediated upregulated and downregulated abiotic stress-associated, unique-expressed orthologs showed lower basal and higher % induction/reduction in expression in the extremophyte (Figs. 4A, 4B).

**Figure 4.**
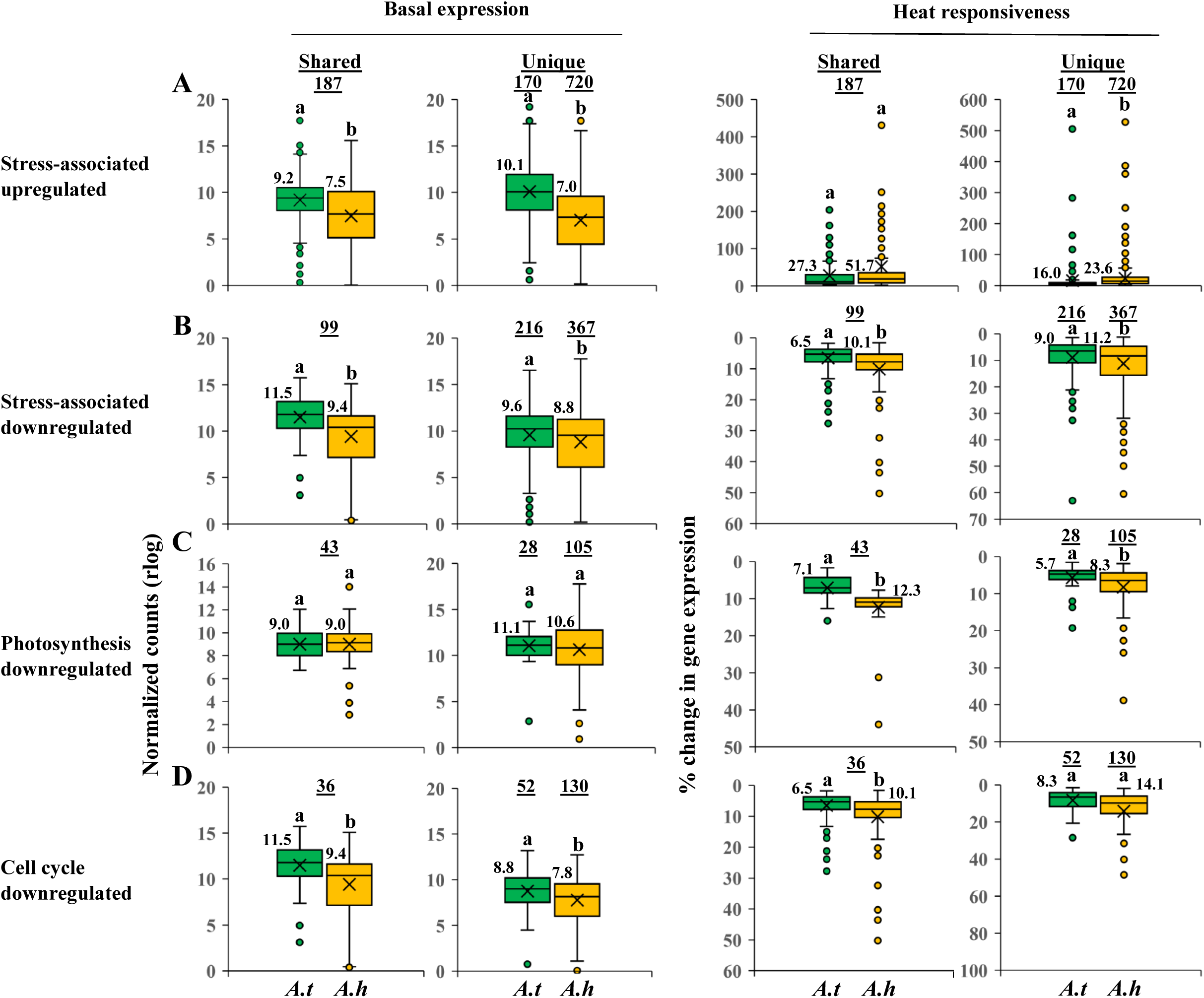
*A. hierochuntica* shared- and unique-expressed orthologs in specific functional groups display lower basal and greater heat-mediated % change in expression than in *A. thaliana*. All genes used in this analysis possess a unique AGI code (putative *A. hierochuntica* orthologs were assigned *A. thaliana* AGI codes). Genes were chosen based on their association with GO terms for their respective categories (Methods S1). Basal expression levels were based on CM conditions. % change in expression from basal level was calculated based on the maximum rlog expression levels of upregulated genes (abiotic stress [A]) or minimum rlog expression levels of downregulated genes (abiotic stress [B], photosynthesis [C], cell cycle [D]) in response to heat stress over the three heat waves. Basal and % change in expression values for all genes in each category are in Dataset S2. For box and whisker plots, the median (thick black line), the mean (cross below the median line) and interquartile range (IQR) of the observed differences are shown. Whiskers indicate the maximum/minimum range. Open circles correspond to extreme observations with values >1.5 times the IQR. Underlined numbers above the circles indicate the number of shared or unique expressed genes. Letters above the circles indicate significant differences at *p* < 0.05 (Student’s *t*-test). Numbers next to boxes are median values. *A.t*, *Arabidopsis thaliana*; *A.h*, *Anastatica hierochuntica*.

Plants actively and early on reduce their growth in response to stress independent of photosynthesis, and this is apparent in a reduction in both cell size and cell elongation that can be linked to downregulation of cell cycle-associated genes (Aguirrezabal et al., 2006; Skirycz et al., 2010; Skirycz et al., 2011; Kazachkova et al., 2013). Subsequently, expression of genes involved in photosynthesis is downregulated under stress (Rizhsky et al., 2002; Zhang et al., 2018a; Huang et al., 2019). We observed that the majority of shared- and unique-expressed orthologs associated with photosynthesis or the cell cycle were downregulated by heat stress in both species (Dataset 2). However, for both shared- and unique-expressed orthologs associated with photosynthesis, *A. hierochuntica* exhibited a similar basal, but greater % reduction in expression than *A. thaliana* (Fig. 4C). Orthologs encoding proteins involved in the cell cycle that possessed shared heat-mediated downregulated expression showed a lower basal and higher % reduction in expression in the extremophyte while unique-expressed cell-cycle orthologs exhibited lower basal expression in *A. hierochuntica* (Fig. 4D).

We next used WGCNA to cluster genes from all conditions into modules with similar expression profiles and detected 22 *A. thaliana* and 21 *A. hierochuntica* co-expression modules (Fig. S3). In both species, two modules clearly covered early heat-induced genes (1.5 h [morning] and 7 h [afternoon] after onset of heat stress) (Fig. 5A; Datasets S4-S8). The morning heat-response modules of both species were enriched in GO biological terms such as “response to heat”, “response to high light intensity” and “response to reactive oxygen species”, (Dataset S9; Methods S1) while the *A. thaliana* and *A. hierochuntica* afternoon heat-response modules were not enriched in any GO-terms. Importantly, both shared- and unique-expressed *A. hierochuntica* genes associated with GO terms for abiotic stress in the early heat-response modules exhibited the same or lower basal expression, and higher heat-mediated % induction of expression than their *A. thaliana* orthologs (Fig. 5B).

**Figure 5.**
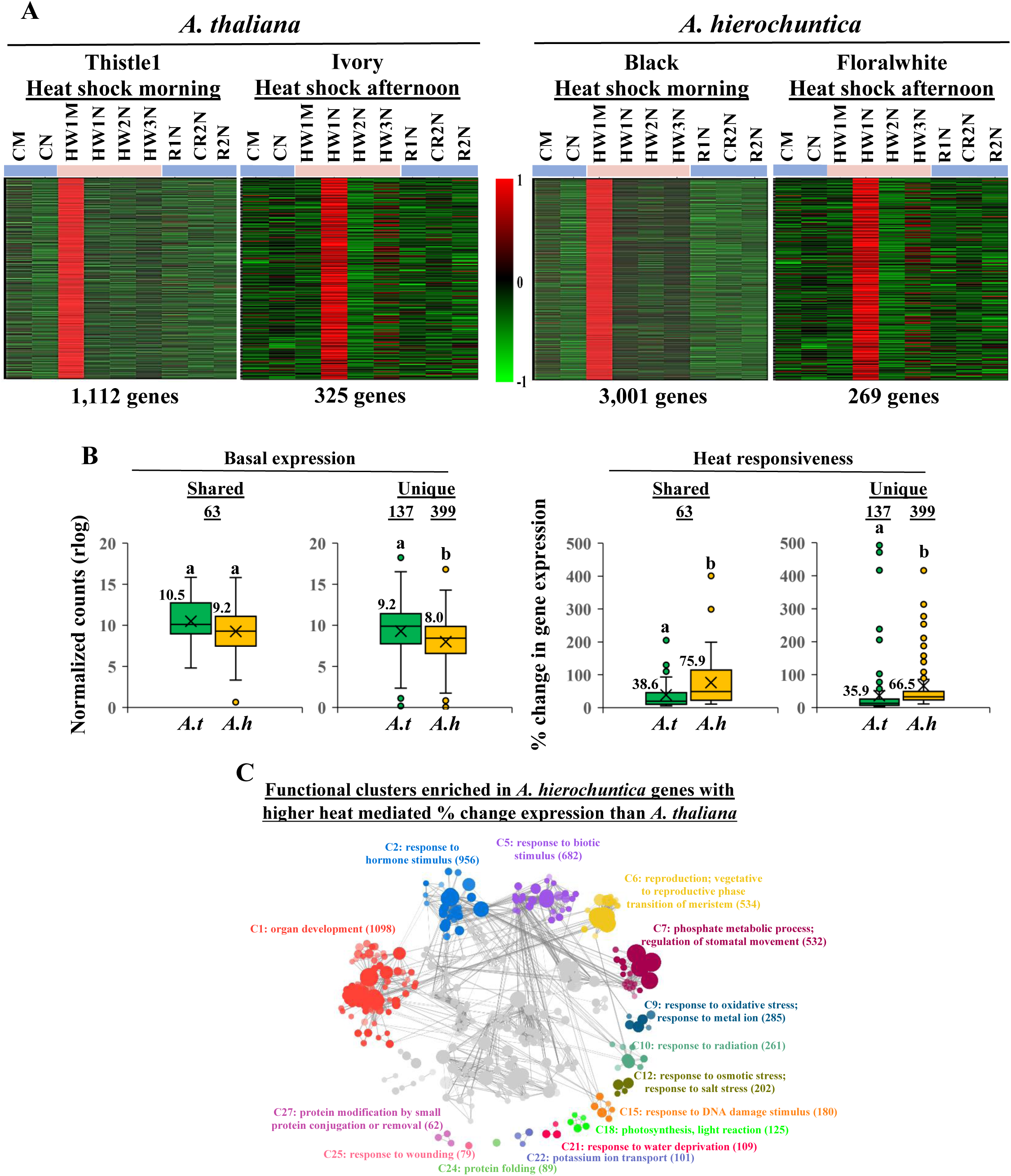
The *A. hierochuntica* early heat response transcriptome displays lower basal and greater heat-mediated % change in expression than in *A. thaliana*. (A), Expression profiles of *A. thaliana* (left two panels) and *A. hierochuntica* (right two panels) morning and afternoon early heat-response modules. These modules were assigned standard color-based names by WGCNA (e.g. Thistle, Ivory etc.; Fig. S3; Datasets S4-S8). Transcript levels were scaled to visualize patterns of expression. The relative intensity of gene expression (red, upregulated; green, downregulated) is shown in the scale bar. Gene expression in each condition represents the average of three biological replicates. The number of genes in each module is shown under the respective module. CM, control morning; CN, control afternoon; HW1M, heat wave 1 morning; HW1N, heat wave 1 afternoon; HW2N, heat wave 2 afternoon; HW3N, heat wave 3 afternoon; R1N, day 1 recovery from heat stress afternoon; CR2N, control plants parallel to the R2N time point afternoon; R2N, day 2 recovery from heat stress afternoon; Blue lines above heat map, control conditions; Pink lines, heat conditions. (B), Expression of orthologs associated with abiotic stress GO terms (Dataset S2; Methods S1) Underlined numbers above the circles indicate the number of shared- or unique-expressed genes. Letters above the circles indicate significant differences at *p* < 0.05 (Student’s T-test). Numbers next to boxes are median values. *A.t*, *Arabidopsis thaliana*; *A.h*, *Anastatica hierochuntica*; (C), Functional clusters enriched among *A. hierochuntica* orthologous genes exhibiting a higher heat-induced % change in expression than *A. thaliana*. For full reactive gene list see Dataset 10. Clustering was performed with the GOMCL tool (https://github.com/Guannan-Wang/GOMCL) (Wang et al., 2020; Methods S1) Clusters are colored differently and labelled with the representative functional term (Dataset 11). Each node represents a gene ontology (GO) term and node size signifies the number of genes in the test set assigned to that functional term; the number of genes in each cluster is in parentheses. The shade of each node represents the *p*-value assigned by the enrichment test (false discovery rate (FDR)-adjusted *p* < 0.05) with darker shades indicating smaller *p*-values. GO-terms sharing > 50% of genes are connected by edges. Only selected clusters are highlighted, the rest are greyed out.

To provide functional support for a more reactive *A. hierochuntica* heat-response transcriptome, we identified 10,653 *A. hierochuntica* genes that exhibited significantly higher heat-mediated % change in expression than their *A. thaliana* orthologs (Dataset S10). These genes were enriched in biological processes related to abiotic stress including oxidative, water, and salt stresses, response to radiation (including genes involved in defense against U/V light), and the response to DNA damage (Fig. 5C, Dataset S11). Additionally, the “protein folding” gene list contained heat shock protein-encoding genes. To validate our gene expression comparisons, we showed (Fig. S4; Methods S1): (i) no significant difference between the species in the proportion of the top 10 most highly expressed genes out of the total transcripts sequenced across all treatments; (ii) similar comparative basal expression results as observed with DEseq2, when we used a new between-species Scale-Based Normalization method (Zhou et al., 2019); (iii) relative and quantitative QPCR analysis confirmation of the RNA-seq fold-change and basal gene expression patterns of selected genes.

Furthermore, we examined the basal expression of 15 orthologous housekeeping genes from both species and found that the average ratio of basal expression of *A. thaliana* to *A. hierochuntica* housekeeping genes was 1.0 ± 0.34 (Fig. S5). Thus, the average lower basal gene expression observed in *A. hierochuntica* compared to *A. thaliana* was not due to lower metabolic activity in the extremophyte.

### Brassicaceae extremophytes possess positively selected genes associated with surviving harsh environments

As a second approach to identifying adaptations to an extremophyte lifestyle, in general, and to desert conditions in particular, we pinpointed positively selected genes (PSGs) that might be indicative of adaptive evolution of stress tolerance. We first used phylogenomics to infer evolutionary relationships between 16 Brassicaceae species including *A. hierochuntica* and representing all major lineages in this family (Dataset S12). *Tarenaya hassleriana* (Cleomaceae) was used as an outgroup. This led to a selection of 13,806 ortholog groups found in 17 taxa. The phylogenomic tree partitioned the species in concordance with their previously assigned lineages (I, II, and III), where *Aethionema arabicum* is considered to belong to a basal clade within the Brassicaceae (Fig. 6A; Franzke et al., 2011; Kiefer et al., 2014). *A. hierochuntica* (*Anastaticeae*) was assigned to LIII (Franzke et al., 2011). It is important to note that *A. hierochuntica* is the single representative species used for LIII due to this lineage being sparsely represented in publicly available genomic databases unlike transcriptomes available for LI and LII species. Thus, to the best of our knowledge, we provide the first substantial genetic resource that enables exploration into adaptive traits that have evolved in a representative lineage III species.

**Figure 6.**
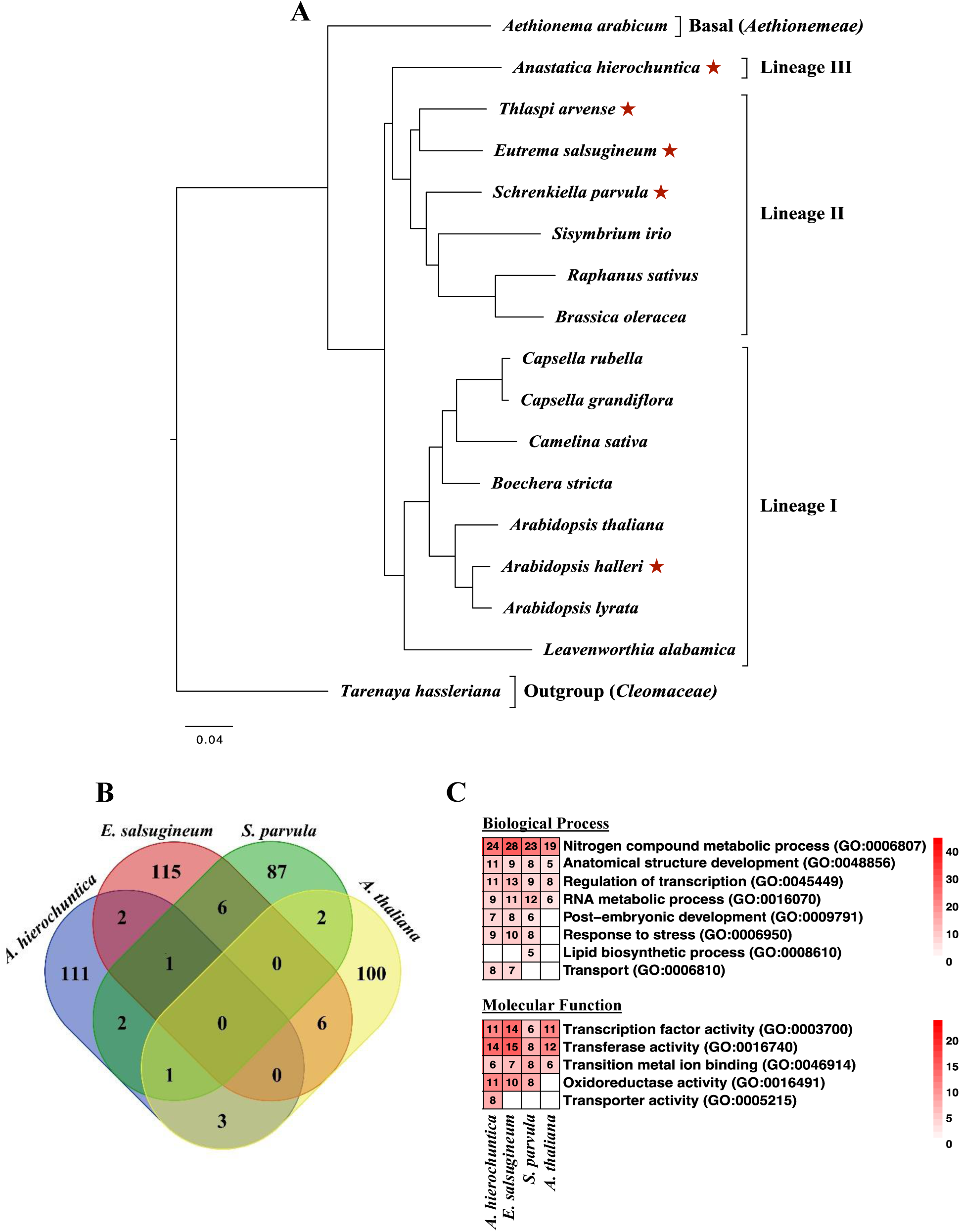
Phylogenomic and comparative positive selection analyses of *A. hierochuntica* and other representative *Brassicaceae* genomes. (A), Maximum-likelihood tree topology based on supermatrix analysis of 13,806 ortholog groups that contain an amino acid sequence from at least four taxa. All nodes are 100% supported by 100 rapid bootstrapping repeats. Red asterisks, extremophyte species; (B), Comparison of the number of PSGs among species. PSGs in each species were identified using the AGALMA-PAML pipeline; (C), Comparative GO-term enrichment analysis of PSGs. The red color intensity corresponds to the number of PSGs assigned with that GO term (the numbers are indicated within the cells). Cells with a white color correspond to GO terms that were not significantly enriched. The *A. thaliana* genome was used as the background gene set and significance (*q*-value < 0.05) of enrichment was assessed via the Fisher’s exact test. For the full list of enriched GO-terms see Fig. S6 and Datasets S18 to S22).

Comparative Brassicaceae transcriptome analysis has revealed that *A. heirochuntica* underwent a mesopolyploid event followed by diploidization (Mandakova et al., 2017). We therefore examined the orthologous groups for any bias towards *A. hierochuntica* using OrthoFinder (Emms and Kelly, 2019). *A. hierochuntica* displayed number of protein coding transcripts, and number and percent of genes present in orthogroups, that were close to the average observed over all 17 species (Fig. S6). These data also suggest that the number of *A. hierochuntica* protein-coding gene models in the curated reference transcriptome is not artificially inflated due to inclusion of a high proportion of alternatively spliced transcripts.

The tree contains five extremophyte species (Fig. 6A, red asterisks): the halophytes *Eutrema salsugineum* and *Schrenkiella parvula* (tolerant to high salinity and multiple other stresses; Kazachkova et al., 2018; Wang et al., 2021a), *Thlaspi arvense* (freezing-tolerant; Sharma et al., 2007; Zhou et al., 2007), *A. hierochuntica* (heat-, salt-, low N-tolerant; Eshel et al., 2017) and *Arabidopsis halleri* (heavy metal hyperaccumulator, semi-alpine conditions; Hanikenne et al., 2008; Honjo and Kudoh, 2019). Therefore, to identify genes under common positive selective pressure in the extremophytes, we used the branch-site model (Yang, 1997; Yang, 2007) to test the external branches (foreground) of the five extremophyte species against all the other branches (background). We then repeated this procedure to test for PSGs in three specific extremophytes - the well-studied halophyte models, *E. salsugineum* and *S. parvula*, and *A hierochuntica* - by labelling each species’ external branch as the foreground. Overall, we identified 194, 120, 130 and 99 genes with a positive selection signal in the “all extremophyte species”, *A. hierochuntica*, *E. salsugineum*, and *S. parvula* runs, respectively (Datasets S13-S16). We also tested *A. thaliana*, as an abiotic stress-sensitive control, and identified 112 PSGs (Dataset S17).

We next examined whether any common genes are under positive selection in the extremophytes or between the extremophytes and stress-sensitive *A. thaliana*. While we could not detect a clear convergence in the use of common positively selected orthologs in the extremophytes (Fig. 6B), the functional attributes shared by those positively selected orthologs in each extremophyte exhibited convergence (Fig. 6C; Fig. S7; Datasets S18 to S22). Notably, orthologs associated with the GO-term “response to stress” [GO:0006950] were highly enriched in the extremophytes suggesting major selective pressure for stress tolerance imposed by their extreme environments.

PSGs from the “all extremophyte species”, supported association with adaptations to harsh environments. For instance, *AKS2*, *MYB52*, *WRKY75*, *ASF1B* and *PHR1*/*UVR2* that have known functions in ABA responses, phosphate starvation, heat stress, and UV-B radiation stress (Table 1 and refs. therein), were among the positively selected group of genes in the extremophytes. Interestingly, *bZIP1* (salt/drought tolerance, and nitrogen signaling) and *APX6* (ROS-scavenging) showed signatures of positive selection unique to *A. hierochuntica* (Table 1), which is highly tolerant to low N and oxidative stresses, and moderately tolerant to salt stress (Eshel et al., 2017). PSGs unique to *S. parvula* included *CAX11*/*CCX5* and *RAB28* that are involved in high-affinity K^+^ uptake and Na^+^ transport, and lithium toxicity, respectively (Table 1; Borrell et al., 2002; Zhang et al., 2011). The pinpointing of these two genes added validity to our positive selection analysis because the native soils of *S. parvula* contain highly toxic levels of Li^+^ and K^+^ (Helvaci et al., 2004; Ozfidan-Konakci et al., 2016), and this species displays extreme tolerance to both Li^+^ and K^+^ toxicity (Oh et al., 2014; Pantha et al., 2021). In contrast to the extremophyte species, PSGs in *A. thaliana* were related to biotic stress responses (Table 1). Of the PSGs in the “all extremophyte species” or *A. hierochuntica*-specific sets, *AKS2*, *bZIP1* and *PHR1/UVR1* expression displayed clear and significantly higher transcript levels in *A. hierochuntica* compared to *A. thaliana* whereas the expression of *APX6* exhibited lower transcript levels in *A. hierochuntica* (Fig. 7A). Exclusively in *A. hierochuntica,* we identified, *CYP71*, *FAS1*, *FBH2*, *SBI1*/*LCMT1*, and *VIP5* as PSGs that are involved in photoperiodic flowering, regulation of meristems, and control of morphology including shoot branching (Table 1). Furthermore, *AhFAS1* expression was highly upregulated by heat stress while *AtFAS1* expression was downregulated (Fig. 7B). *AhSBI1/LCMT1* expression was unaffected by heat stress whereas *AtSB1/LCMT1* expression was highly upregulated by heat. Moreover, *AhSBI1/LCMT1* transcript levels were lower than *AtSBI1/LCMT1* over all time points. Notably, genes involved in organ development and flowering time were more reactive to heat in *A. hierochuntica* than in *A. thaliana* (Fig. 5C). Considering that *A. hierochuntica* ontogeny is very different from *A. thaliana*, *E. salsugineum* and *S. parvula* – it exhibits a multi-branched sympodial shoot structure supporting multiple axillary inflorescences that flower independent of day length (Fig. 1B; Gutterman, 1998; Eshel et al., 2017) - positive selection of these genes could indicate an important adaptation to the desert environment.

**Figure 7.**
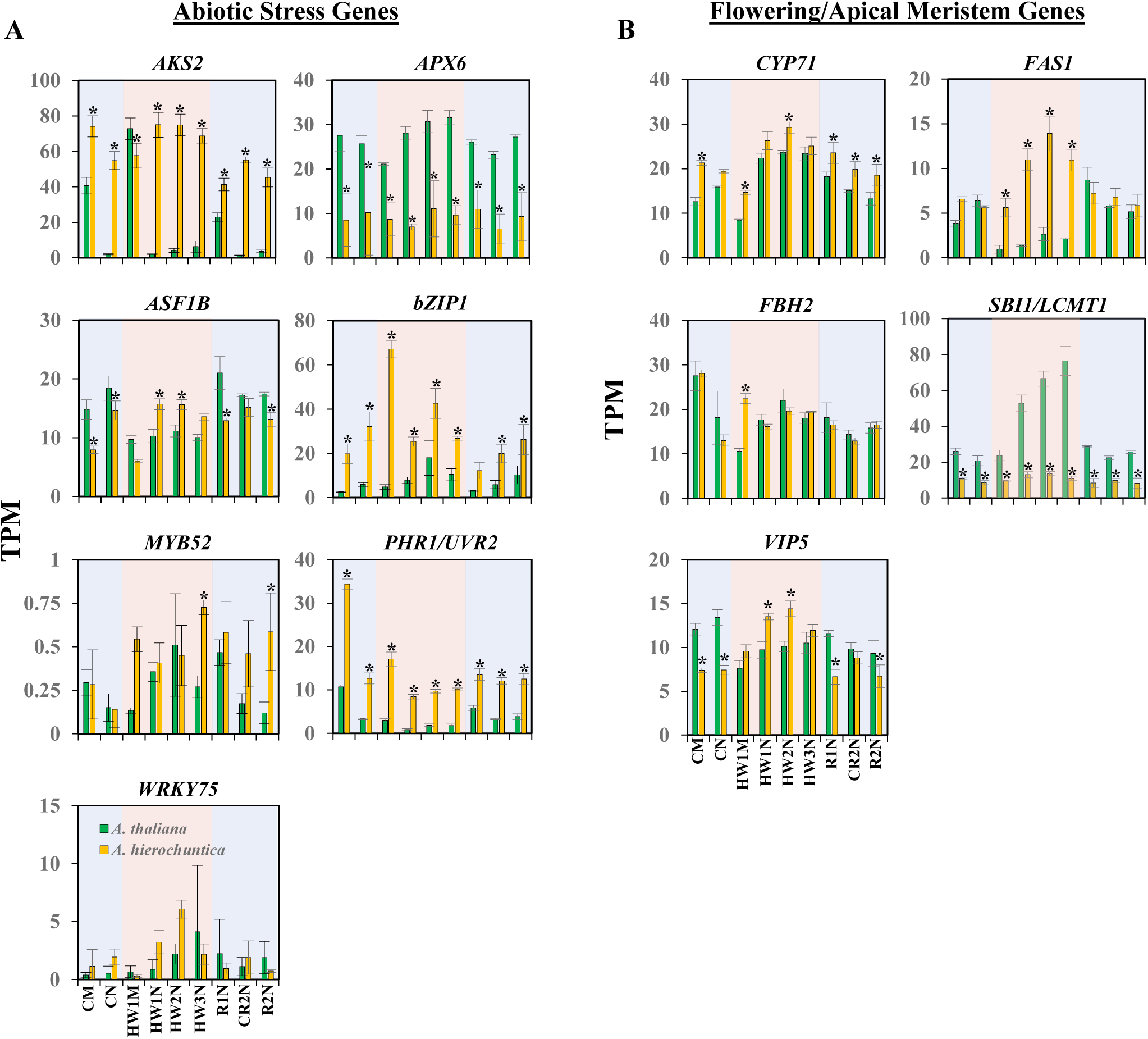
Expression of PSGs in response to heat stress. Gene expression was assessed by RNA-seq transcriptome analysis of *A. thaliana* and *A. hierochuntica* plants grown under control conditions or exposed to heat stress (see Fig. 2A for experimental design). Expression is expressed as transcripts per kilobase million (TPM) normalized gene expression. (A), PSGs from the “all extremophyte species” and *A. hierochuntica* analyses that are associated with abiotic stress responses (Table 1); (B), *A. hierochuntica* PSGs that function in photoperiodic flowering, regulation of meristems, and control of morphology (Table 1). Data are mean ± S.D. (n = 3) Asterisks indicate significant difference at *p* < 0.05 between *A. thaliana* and *A. hierochuntica* at the same time point and condition (Student’s *t*-test). CM, control morning; CN, control afternoon; HW1M, heat wave 1 morning; HW1N, heat wave 1 afternoon; HW2N, heat wave 2 afternoon; HW3N, heat wave 3 afternoon; R1N, day 1 recovery from heat stress afternoon; CR2N, control plants parallel to the R2N time point afternoon; R2N, day 2 recovery from heat stress afternoon; Blue shading, control conditions; Pink shading, heat conditions.

**Table 1.** Positively selected genes**^a^**, with a potential role in adaptation to extreme environments. Selected genes from five CODEML branch-site model analyses are indicated based on their *A. thaliana* ortholog identifier. TF, Transcription Factor.

| Positively selected gene | Function | References |
| --- | --- | --- |
| **All extremophyte species** (*A. hierochuntica*, *E. salsugineum*, *S. parvula*, *T. arvense* and *A. halleri*) |  |  |
| *AKS2* (At1g05805) | TF; facilitates stomatal opening, ABA response | Takahashi et. al. 2013 |
| *ASF1B* (At5g38110) | Histone H3/H4 chaperone; repair of UV-B-induced DNA damage, basal and acquired thermotolerance | Lario et al., 2013; Nie et al., 2014; Weng et al., 2014 |
| *MYB52* (At1g17950) | TF; ABA response, drought tolerance, involved in the regulation of secondary wall formation, seed mucilage | Park et al., 2011; Cassan-Wang et al., 2013; Shi et al., 2018 |
| *PHR1/UVR2* (At1g12370) | Photolyase enzyme; repair of UV-B-induced DNA damage | Ahmad et al., 1997; Landry et al., 1997; Jiang et al., 1997 |
| *WRKY75* (At5g13080) | TF; Pi starvation, root development, GA-mediated flowering, defense response | Devaiah et al., 2007; Velasco et al., 2016; Guo et al., 2017; Zhang et al., 2018b |
| ***A. hierochuntica*** |  |  |
| *APX6* (At4g32320) | Hydrogen peroxide-scavenging enzyme; alleviation of ROS damage | Chen et al., 2014 |
| *bZIP1* (At5g49450) | TF, light and nitrogen sensing, salt and drought tolerance | Obertello et al., 2010; Sun et al., 2012; Para et al., 2014 |
| *CYP71* (At3g44600) | Cyclophilin; silencing of homeotic genes; meristem development, interacts with FAS1 and the floral repressor LHP1 | Li et al., 2007; Li and Luan, 2011 |
| *FAS1* (At1g65470) | Subunit of CaF-1; organization of apical meristems, cellular differentiation, DNA repair | Leyser & Furner, 1992; Kaya et al., 2001; Hisanaga et al., 2013 |
| *FBH2* (At4g09180) | TF; photoperiodic flowering | Ito et al., 2012 |
| *SBI1/LCMT1* (At1g02100) | Leucine carboxylmethyltransferase; brassinosteroid signaling; flowering, stress responses | Di Rubbo et al., 2011; Wu et al., 2011; Creighton et al., 2017 |
| *VIP5* (At1g61040) | PAF1c component; activates floral repressors and photoperiodic pathway regulators. regulation of N uptake | Oh et al., 2004; Yu and Michaels, 2010; Crevillen & Dean, 2011; Widiez et al., 2011; Lu et al., 2017 |
| ***E. salsugineum*** |  |  |
| *ATCES1/ACER* (At4g22330) | Alkaline ceramidase; sphingolipid homeostasis, disease resistance, salt tolerance | Wu et al., 2015 |
| *GRXS13* (At1g03850) | Glutaredoxin; Chilling and photooxidative stress tolerance | Laporte et al., 2012; Hu et al., 2015 |
| *NCA1* (At3g54360) | Chaperone; regulates catalase 2 (ROS-scavenging enzyme) activity, salt, cold, high pH stresses | Li et al., 2015 |
| *PSRP2* (At3g52150) | Plastid-specific ribosomal protein; RNA chaperone activity, negative regulator of seed germination under abiotic stress | Xu et al., 2013 |
| *SLK2* (At5g62090) | Transcriptional adaptor; embryogenesis, organ development, repression of stress-responsive gene transcription | Bao et al., 2010; Lee et al., 2014; Shrestha et al., 2014 |
| ***S. parvula*** |  |  |
| *Atrab28* (At1g03120) | LEA protein; Li<sup>+</sup> tolerance | Borrell et al., 2002 |
| *CAX11/CCX5* (At1g08960) | Cation calcium exchanger; K<sup>+</sup> uptake, Na<sup>+</sup> transport in yeast | Zhang et al., 2011 |
| *PER1* (At1g48130) | Peroxioredoxin; ROS scavenging, enhances primary seed dormancy | Chen et al., 2020 |
| ***A. thaliana*** |  |  |
| *ATG6* (At3g61710) | AuTophGy-related protein; autophagy, pathogen defense | Patel & Dinesh-Kumar, 2008 |
| *ATL2* (At3g16720) | RING-H2 zinc-finger protein; pathogen defense | Serrano & Guzmán, 2004 |
| *ERDJ3B* (At3g62600) | ER-localized DNAJ chaperone; anther development under heat stress, pathogen defense | Nekrasov et al., 2009; Yamamoto et al., 2020 |
| *RST1* (At3g27670) | ARM-repeat protein; RNA exosome cofactor, vacuolar trafficking, cuticular wax production, embryo development, pathogen defense | Chen et al., 2005; Mang et al., 2009; Lange et al., 2019; Zhao et al., 2019; |
| *XLG2* (At4g34390) | Heterotrimeric G protein; pathogen defense | Liang et al., 2016 |

<sup>a</sup>For log-likelihood values of the alternative and null models, log-likelihood ratio tests and *p*-values, see Datasets S13 to S17.

## Discussion

### The *A. hierochuntica* transcriptome does not exist in a heat “stress-ready” state and is more reactive to heat stress than *A. thaliana*

Our finding that *A. thaliana* and *A. hierochuntica* exhibit similar transcriptome adjustment in response to heat stress and during recovery (Figs. 2E, 2F) distinguishes *A. hierochuntica* from other extremophyte relatives. The extent of transcriptomic, proteomic and metabolic adjustment in response to ionic stress in the halophytes *E. salsugineum* and *S. parvula*, is much lower than in *A. thaliana* (Taji et al., 2004; Gong et al., 2005; Lugan et al., 2010; Pang et al., 2010; Oh et al., 2014; Vera-Estrella et al., 2014; Wang et al., 2021a). This lower adjustment reflects their “stress-ready” state whereby transcript and metabolite accumulation that is induced or repressed in *A. thaliana* in response to ionic stress, is constitutively high or low, respectively, in the halophytes (Gong et al., 2005; Lugan et al., 2010; Kazachkova et al., 2013; Oh et al., 2014; Wang et al., 2021a). A “stress-ready” transcriptome is exemplified in *S. parvula* where basal expression of over 1,000 “stress-ready” orthologs matches the post-boron stress expression levels observed in *A. thaliana* (Wang et al., 2021a). In contrast, the great majority of *A. hierochuntica* and *A. thaliana* orthologs exhibit a shared response mode (Fig. 3A). Furthermore, many stress-related *A. hierochuntica* genes show lower basal and/or higher fold-change gene expression compared to *A. thaliana* (Figs. 3, 4, 5). Indeed, almost one third of *A. hierochuntica* genes display a higher heat-mediated fold-change in expression compared to *A. thaliana* and are enriched in abiotic stress-associated functions (Fig. 5; Datasets 10, 11). Taken together, our findings support a paradigm whereby the *A. hierochuntica* transcriptome is more reactive to heat stress than *A. thaliana*.

One reason for the difference in the transcriptome responses of *A. hierochuntica* vs. its halophytic relatives relates to the type of stress each species faces. *E. salsugineum* and *S. parvula* habitats possess levels of ions such as Na^+^ and BO ^3-^ that are toxic to the majority of plant species and the two halophytes are constantly exposed to ionic stress throughout their life cycle. This situation might have led to the evolution of a stress-associated transcriptome that is continuously “switched-on”. Conversely, *A. hierochuntica* is generally exposed to heat stress later in its life cycle and on a diurnal basis thus favoring a more reactive transcriptome.

A second reason is that *A. hierochuntica* thrives in an environment with seasonal temperatures ranging from -3.6 to 46.8 °C (Eshel et al., 2017), with diurnal maximum variations exceeding 18 °C (Israeli Meteorological Services). On the other hand, *E. salsugineum* can be found in locations such as China’s Shandong peninsula where temperatures range from -5 °C in the winter to 32 °C in the summer and with diurnal temperature differences rarely exceeding 10 °C (Guedes et al., 2015). Furthermore, diurnal temperature differences in the cold, spring-growing period of *E. salsugineum* are likely more moderate than those experienced by *A. hierochuntica* during the warm Negev desert spring. Thus, evolution of a flexible transcriptome that confers a strong reaction to extreme diurnal temperature fluctuations could be advantageous for adaptation to a desert environment. Moreover, a transcriptome with globally lower basal expression levels would require less energy to be expended in the low nutrient desert environment.

### Brassicaceae extremophytes possess common PSGs that are indicative of adaptation to harsh environments

Extremophytes are present in all three Brassicaceae lineages (Fig. 6A; Franzke et al., 2011) illustrating that adaptation to stressful habitats has occurred independently, multiple times within the Brassicaceae and is indicative of convergent evolution (Kazachkova et al., 2018; Birkeland et al., 2020). Consistent with this notion, we identified a set of 194 genes under positive selection across the five extremophyte species that could commonly contribute to plant adaptation to extreme environments. Other studies with extremophyte Brassicaceae have also detected signatures of positive selection in genes that function in stress tolerance (Zhou et al., 2009; Jarvis et al., 2014; Birkeland et al., 2020). For instance, positive selection of stress-associated genes was detected in three Arctic Brassicaceae species (Birkeland et al., 2020). Similar to our findings (Figs. 6B, 6C, S6) there was little overlap of PSGs between the Arctic extremophytes but considerable overlap in functional pathways. Taken together, these data do not support adaptive molecular convergence but rather indicate evolution of similar adaptations via distinct evolutionary pathways.

Among the genes under positive selection across the five extremophyte species in the current study, we identified two genes encoding ABA-responsive transcription factors (TFs), AKS2 and MYB52 (Table 1), illustrating the importance of the ABA response networks in adaptive evolution of stress tolerance (Xia et al., 2010; Fischer et al, 2011; Bondel et al., 2018). In particular, the bHLH TF, ABA-RESPONSIVE KINASE SUBSTRATE (AKS) 2 activates transcription of K^+^ channels in stomata guard cells in an ABA-dependent manner thereby enhancing stomatal opening (Takahashi et al., 2013). It is intriguing that a regulator of stomatal aperture has undergone positive selection across the extremophytes because alterations in stomatal aperture is a crucial early response to multiple abiotic stresses (Brugnoli and Lauteri, 1991; Chaves et al., 2009; Stepien and Johnson, 2009; Devireddy et al., 2020). Thus, positive selection of non-synonymous amino acid changes in the coding region of *AKS2*, (plus differences in heat-mediated regulation of *A. thaliana* and *A. hierochuntica AKS2* expression (Fig. 7A)) suggest that this gene may have been naturally selected for survival in extreme environments.

*WRKY75* was also positively selected across the extremophytes (Table 1). In *A. thaliana*, this TF regulates the expression of several key phosphate starvation-induced genes (Devaiah et al., 2007). Extremophytes often exist on soils with low Pi availability (Thompson et al., 2006; Holzapfel, 2008; Guevara et al., 2012). For instance, the *E. salsugineum* Yukon ecotype grows in the low Pi soils of the Yukon region in Canada (Guevara et al., 2012) and is highly tolerant to Pi deficiency compared to *A. thaliana*. This tolerance is associated with higher basal expression of several Pi starvation genes including *WRKY75* (Velasco et al., 2016). Both *E. salsugineum* and *A. hierochuntica* exhibit significantly higher basal levels of Pi than *A. thaliana* (Gong et al., 2005; Kazachkova et al., 2013; Velasco et al., 2016; Eshel et al., 2017). Thus, positive selection of *WRKY75* across the five extremophyte plants, and differential expression of *A. thaliana* and *E. salsugineum WRKY75* suggests that selection for more efficient extraction of soil Pi is a common evolutionary adaptation to extreme environments.

Extremophytes are often exposed to UV-B radiation that can cause direct damage to DNA (Kimura and Sakaguchi, 2006). *PHOTOLYASE1*/*UV-RESISTANCE2* (*PHR1*/*UVR2*) and *ANTI-SILENCING FUNCTION 1b* (*ASF1B*) that are crucial for repairing UV-B-induced DNA damage were positively selected across the five extremophytes (Table 1; Ahmad et al., 1997; Landry et al., 1997; Jiang et al., 1997; Lario et al., 2013; Nie et al., 2014). *PHR1*/*UVR2* is also the major mechanism maintaining transgenerational genome stability in *A. thaliana* continuously exposed to UV-B (Willing et al., 2016) while *ASF1B* is also involved in the regulation of basal and acquired thermotolerance (Weng et al., 2014). Additionally, *PHR1*/*UVR2* displays higher basal expression in *A. hierochuntica* compared to *A. thaliana*, and while heat leads to downregulation of the gene in both species, expression is reduced to a lesser extent in the extremophyte. (Fig. 7A).

Collectively then, our data suggest common selective pressures in extremophyte plants that target key components in stomatal opening, nutrient acquisition, and UV-B-induced DNA repair. On the other hand, we found that *A. thaliana* PSGs were principally involved in defense against pathogens (Table 1). This supports the hypothesis that because *A. thaliana* evolved in temperate regions where pathogen density is relatively high compared to extremophyte habitats, it encountered greater evolutionary pressures for adaptation to biotic stresses (Oh et al., 2014).

### *A. hierochuntica* PSGs suggest adaptive evolution for an opportunistic desert lifestyle

We pinpointed a number of PSGs specifically in *A. hierochuntica* indicating adaptation to the desert environment (Table 1; Fig. 7B). Intriguingly, several of these genes function in *A. thaliana* in the transition from vegetative to reproductive growth and meristem development: (i) *VERNALIZATION INDEPENDENCE VIP5* enhances transcription of the floral repressor *FLOWERING LOCUS C* (*FLC*) gene and other *MADS AFFECTING FLOWERING* (*MAF*) gene family members (Oh et al., 2004; Yu and Michaels, 2010; Crevillen and Dean, 2011; Lu et al., 2017) and *A. thaliana vip5* mutants exhibit early flowering; (ii) FLOWERING BHLH 2 (FBH2) activates transcription of the *CONSTANS* gene, a central regulator of photoperiodic flowering. *FBH2* overexpression causes photoperiod-independent early flowering (Ito et al., 2012); (iii) *FASCIATA1* (*FAS1*) appears to function in the organization of shoot and root apical meristems, and in cellular differentiation (Kaya et al., 2001; Exner et al., 2006). Mutations in *fas1* cause stem fasciation, abnormal leaf and flower morphology, and defects in the organization of apical meristems (Leyser and Furner, 1992; Kaya et al., 2001); (iv) *CYP71* plays a critical role in regulating meristem development, including the floral meristem (Li et al., 2007). Furthermore, CYP71 physically interacts with FAS1 thereby targeting FAS1 to the *KNAT1* locus (Li and Luan, 2011). KNAT1 is essential for maintenance of apical meristems (Hake et al., 2004). In addition, CYP71 interacts with LIKE HETEROCHROMATIN PROTEIN 1 (LHP1), which is involved in repressing expression of flowering time and floral identity genes (Gaudin et al., 2001; Kotake et al., 2003). Thus, *lhp1* mutations cause strong early flowering; (v) *SUPPRESSOR OF BRI1 (SBI1)*/*LEUCINE CARBOXYLMETHYLTRANSFERASE (LCMT1)* regulates components of the brassinosteroid signaling pathway (Di Rubbo et al., 2011; Wu et al., 2011) and the *sbi1/lcmt* mutant is early flowering in both long and short days consistent with the role of brassinosteroids in flowering (Li and He, 2010; Nolan et al., 2020).

The discovery of positively selected flowering and meristem development genes specifically in *A. hierochuntica* is consistent with its very different developmental program compared to many other Brassicaceae including the additional four extremophyte plants included in our analysis. *A. hierochuntica* does not display the distinctive transition from the vegetative rosette leaf stage to the reproductive bolting stage, which is accelerated in long-day conditions (Pouteau and Albertini, 2009; Song et al., 2013). Instead, regardless of photoperiod, the shoot repeatedly bifurcates from the four true-leaf stage onwards, developing an axillary inflorescence at each branch point thereby leading to a multi-branched shoot morphology (Fig. 1B, panel (i); Eshel et al., 2017). Most interestingly, mutation in the *A. thaliana FAS1* gene (whose *A. hierochuntica* ortholog is under positive selection) can induce stem bifurcation and enlargement (Leyser and Furner, 1992). The shoot bifurcation, multi-branch, photoperiod-insensitive, early flowering traits could maximize fitness in the unpredictable desert environment where plants need to ensure development of seeds but might not survive until a critical day length induces flowering. This idea is supported by our observations of *A. hierochuntica* populations in the Dead Sea valley of Israel, where tiny dead plants that have still managed to produce a few seeds can be seen alongside much larger plants presumably from a year with higher rainfall (Fig. 1B).

In conclusion, we have shown that *A. hierochuntica* possesses a more reactive heat-response transcriptome, and stress-related genes that have undergone positive selection. Genes that could be associated with its multi-branch, early flowering phenotype also exhibit signatures of positive selection. Together, these evolutionary adaptations could allow survival in a hot desert environment with unpredictable precipitation. Our study furthermore provides rich gene sets that will facilitate comparative and functional genomics studies to reveal additional molecular mechanisms for plant tolerance to heat stress in a desert habitat.

## Acknowledgements

We would like to dedicate this paper to Dr. Dirk Hincha and Prof. Hillel Fromm, who passed away in 2020. Great plant scientists, colleagues and friends. They will be sorely missed. We thank Ruth Shaked and Beery Yaakov for their dedicated technical support. We are also grateful to Patrick Barko from the University of Illinois Urbana-Champaign for excellent aid with WGCNA. Our appreciation to Alvaro Hernandez and all the team at the Roy J. Carver Biotechnology Center, University of Illinois Urbana-Champaign for superb sequencing services. We are grateful to Julie Galbraith (Glasgow Polyomics) for preparing libraries and carrying out the Illumina RNA-sequencing for the reference transcriptome. This work was supported by the Goldinger Trust Jewish Fund for the Future, the Koshland Foundation for Support of Interdisciplinary Research In Combatting Desertification, and the I-CORE Program of the Planning and Budgeting Committee to S.B., National Science Foundation award MCB 1616827, Next-Generation BioGreen21 Program of Republic of Korea (PJ01317301).to M.D., NSF-BSF-IOS-EDGE 1923589/2019610 to M.D. and S.B., and the Biotechnology and Biological Sciences Research Council grant (BBSRC; BB/R019894/1) to AA and PH. G.E. was supported by an Israel President Fellowship for Excellence and Scientific Innovation award, N.D was supported by a Ben-Gurion University, Kreitman School for Advanced Research Studies High-tech, Bio-tech and Chemo-tech award, and G.W. was supported by an Economic Development Assistantship award from Louisiana State University, The authors also acknowledge the LSU High Performance Computing Services, and the BGU Bioinformatics Core Facility for providing computational resources needed for data analyses.

## Author contributions

S.B. conceptualized and supervised the overall project; GE and ND performed the main analyses; GW contributed to transcriptome response assessment and performed the GOMCL analysis; GE, D-HO, MG, MD, and VC-C contributed to assembly of the *A. hierochuntica* reference transcriptome. MD assisted with, and GE performed the positive selection analysis; YK and MAO contributed to plant growth, preparation of samples and transcriptome response validation; AA and PH designed, supervised and analysed the Illumina RNA-sequencing experiment for the reference transcriptome at Glasgow Polyomics; AM-C contributed to the WGCNA; GE, ND, GW, SB and MD contributed to data interpretation, GE, ND, and SB wrote the article. MD and AA critically revised and approved the final manuscript.

## Data availability

Reference transcriptome and RNA-seq reads as well as the fully assembled transcriptome are openly available via the NCBI SRA and TSA databases under BioProject PRJNA731383. Other main data that supports the findings of this study are available in the main text and Supporting Information of this article.

## Supporting Information

**Fig. S1.** Transcriptome sequencing and hybrid assembly workflow.

**Fig. S2.** An example of a three-day heat wave event in the Dead Sea valley during April 2008.

**Fig. S3.** Clustering dendrogram of module eigenvalues for *A. thaliana* and *A. hierochuntica* transcriptome profiles under and heat stress conditions.

**Fig. S4.** Validation of “between species” RNA-seq analysis.

**Fig. S5.** Basal (control) expression of 15 orthologous *A. thaliana* and *A. hierochuntica* housekeeping genes.

**Fig. S6.** Number of protein-coding transcripts and comparative ortholog group composition for species used to detect positively selected genes

**Fig. S7.** GO-term enrichment analysis of positively selected genes.

**Dataset S1.** *A. hierochuntica* transcriptome functional annotation.

**Dataset S2.** *A. thaliana* and *A. hierochuntica* raw read data plus GO-terms, and DEGs.

**Dataset S3.** Assignment of ortholog pairs modes of expression (log_2_ fold-change data and WGCNA modules).

**Dataset S4.** Early heat-response modules (rlog expression data).

**Dataset S5.** *A. thaliana* early heat module (Thistle1) gene list.

**Dataset S6.** *A. thaliana* late heat module (Ivory) gene list.

**Dataset S7.** *A. hierochuntica* early heat module (Black) gene list.

**Dataset S8.** *A. hierochuntica* early heat module (Floralwhite) gene list.

**Dataset S9.** GO-term enrichment of *A. thaliana* and *A. hierochuntica* morning heat-response modules. **Dataset S10.** Difference in percent change orthologous gene expression (based on TPM) between *A. heirochuntica* and *A. thaliana*.

**Dataset S11.** GO-term enrichment of genes that are more responsive to heat in *A. hierochuntica* (see Fig. 5C).

**Dataset S12.** Species included in the custom *Brassicaceae* CDS database for the *Brassicaceae* phylogenomic analysis.

**Dataset S13.** CODEML positive selected genes (*q*<0.05) in the all extremophyte species run with the branch-site model.

**Dataset S14.** CODEML positive selected genes (*q*<0.05) in *A. hierochuntica* run with the branch-site model.

**Dataset S15.** CODEML positive selected genes (*q*<0.05) in *E. salsugineum* run with the branch-site model.

**Dataset S16.** CODEML positive selected genes (*q*<0.05) in *S. parvula* run with the branch-site model. **Dataset S17**. CODEML positive selected genes (*q*<0.05) in *A. thaliana* run with the branch-site model. **Dataset S18.** GO-terms over-represented in the all extremophyte species 194 positive selected genes.

**Dataset S19.** GO-terms over-represented in *A. hierochuntica* 120 positive selected genes. **Dataset S20.** GO-terms over-represented in *E. salsugineum* 131 positive selected genes. **Dataset S21.** GO-terms over-represented in *S. parvula* 99 positive selected genes.

**Dataset S22.** GO-terms over-represented in *A. thaliana* 112 positive selected genes.

**Dataset S23.** 109 most conserved orthologs (≥ 98% query coverage, ≥ 99% nucleotide identity, ≥ BLAST value of e-100) between *A. thaliana* and *A. hierochuntica*.

**Dataset S24.** PCR primers used in this study.

**Methods S1**

## Notes

### Competing Interest Statement

The authors have declared no competing interest.

### Summary of Updates

Text corrections have been added for clarification. A new analysis had been added to the text including a new Supplementary Figure S6 to further validate the positive selection analysis.

