## Supplemental Figures for "Positive Selection and Heat-Response Transcriptomes Reveal Adaptive Features of the Brassicaceae Desert Model, *Anastatica hierochuntica*"

### Supplementary Figures

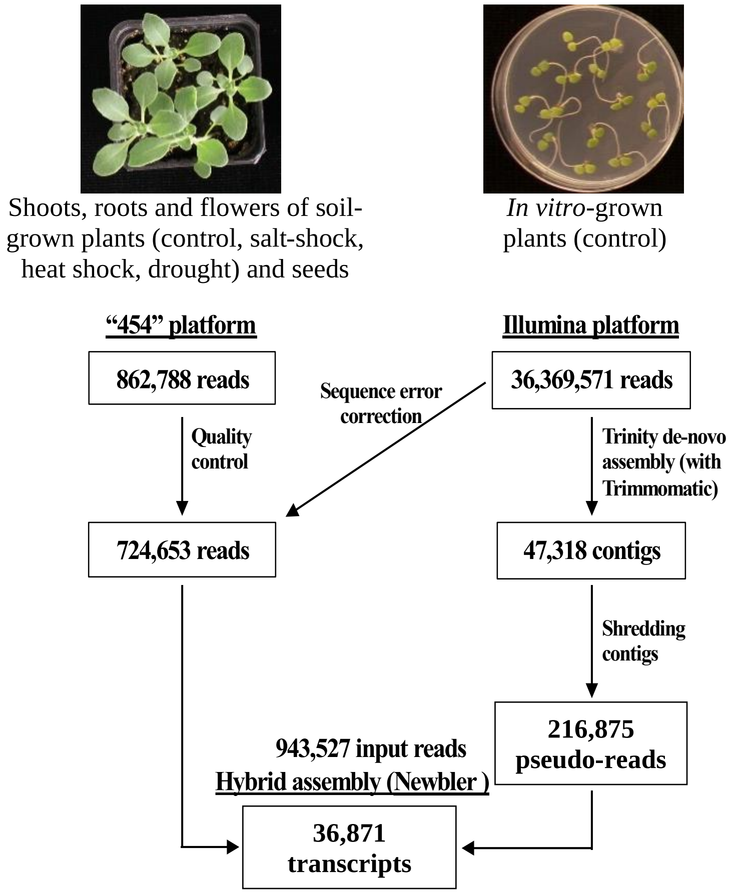


**Figure S1. Transcriptome sequencing and hybrid assembly workflow.** RNA was extracted from plate-grown *A. hierochuntica* seedlings under control conditions (Illumina sequencing), or from soil-grown plants under control or various abiotic stress conditions, and imbibed seeds (454 sequencing). To assemble both short- and long-read sequences together, a hybrid approach was taken. The Illumnia reads were first assembled into long contig sequences, using the Trinity assembler, and then shredded into 700 bp long, overlapping (at least 200 bp overlap) pseudo-reads that were reassembled together with the 454 reads using the Newbler assembler.

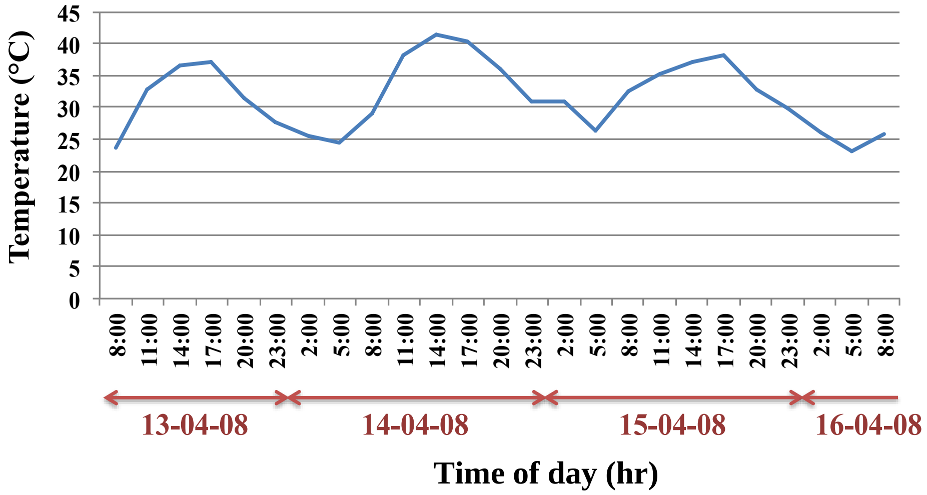


**Figure S2. An example of diurnal temperatures over three days in the Dead Sea valley during April 2008.** Data were obtained from the Israel Meteorological Service (http://www.ims.gov.il/IMSEng).

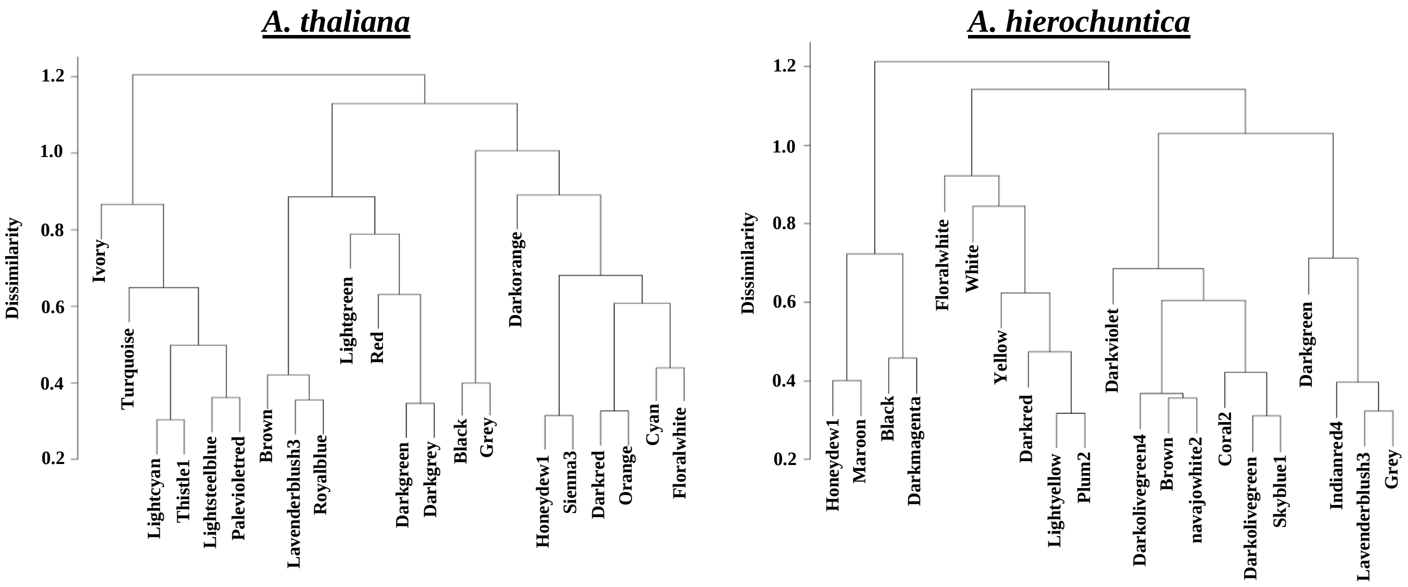


**Figure S3. Clustering dendrogram of module eigenvalues for *A. thaliana* and *A. hierochuntica* transcriptome profiles under and heat stress conditions**. Dendrograms were generated using the WGCNA package with module size minimum set at 50 genes. Dendrograms are presented after merging of modules with a cut off of 30% dissimilarity (70% similarity). These modules were assigned standard color-based names by WGCNA. Identical module names between *A. thaliana* and *A. hierochuntica* do not indicate similarity in function, expression profile or shared genes.

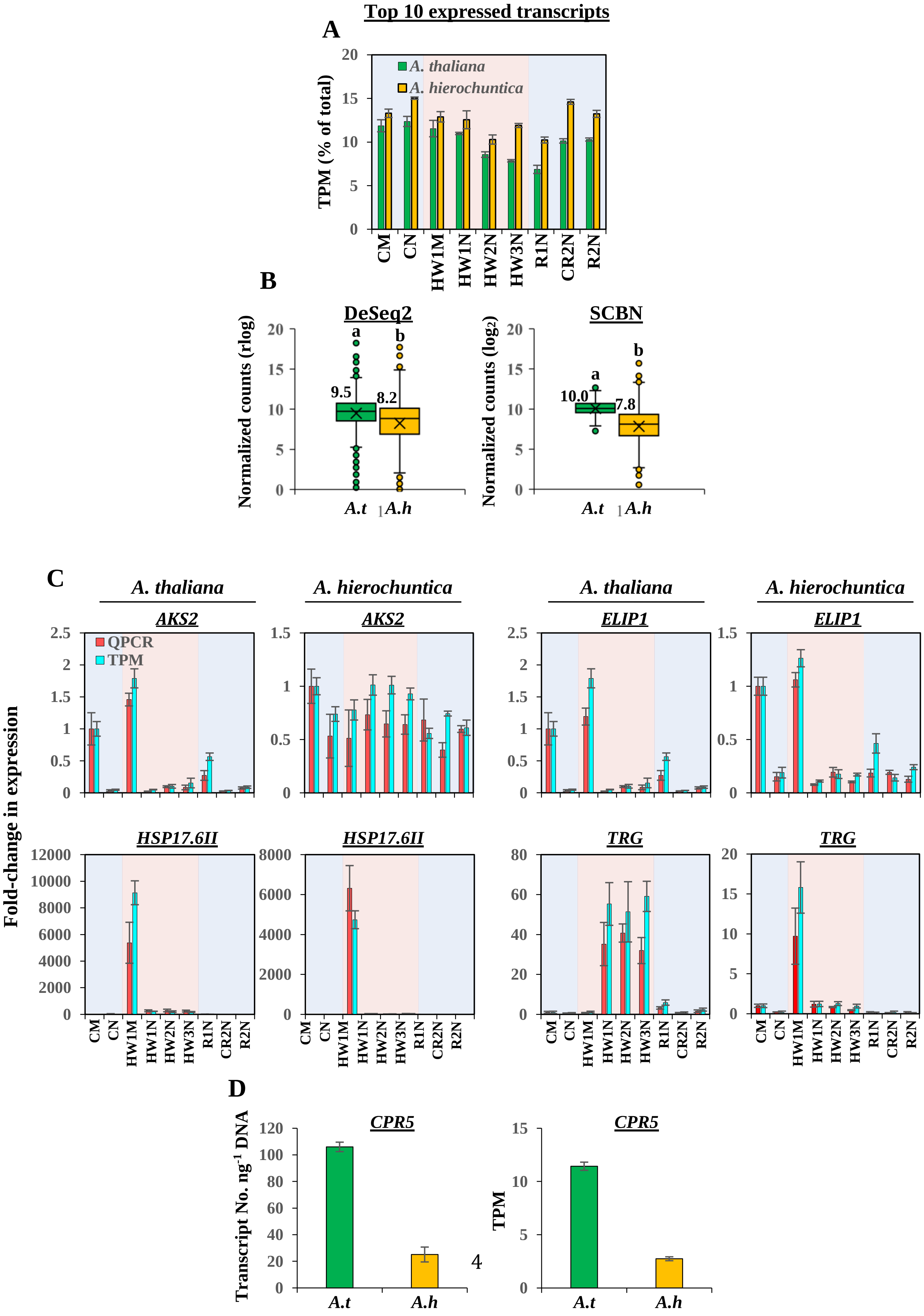
Top 10 expressed transcripts
A
TPM (% of total)
A. thaliana
A. hierochuntica
CM
CN
HW1M
HW1N
HW2N
HW3N
R1N
CR2N
R2N
B
DeSeq2
SCBN
Normalized counts (rlog)
Normalized counts ($log_2$)
a
b
9.5
8.2
10.0
7.8
A.t
A.h
C
A. thaliana
A. hierochuntica
AKS2
ELIP1
HSP17.6II
TRG
QPCR
TPM
Fold-change in expression
D
CPR5
Transcript No. $ng^{-1}$ DNA
TPM
A.t
A.h

**Figure S4. Validation of "between species" RNA-seq analysis.** (A), Top 10 expressed transcripts as a percentage of all expressed transcripts; (B), Comparison of basal expression of control samples using DeSeq2 rlog normalization or the between species Scale-Based Normalization (SCBN) method (Zhou et al., 2019). Numbers next to boxes are median values. Letters above the circles indicate significant differences at $p < 0.05$ (Wilcoxon rank sum test); (C), Relative QPCR expression of selected *A. thaliana* and *A. hierochuntica* genes. Gene expression was determined according to the $2^{-\Delta\Delta C}$T method (Livak and Schmittgen, 2001) using *eIF4A1* from each species as a reference gene. Expression was normalized to the expression level in the control morning sample, which was assigned a value of 1. Data are mean ± SD ($n$ = 3 to 4) and are representative of two independent experiments; (D), Comparison of the basal (control) expression levels of *CPR5* estimated by RNA-seq or absolute QPCR quantification of transcript copy number. Absolute quantification was performed using a fivefold serial dilution of gel-purified *CPR5* and *eIF4A1* (reference gene) QPCR products to create a standard curve. CM, control morning; CN, control afternoon; HW1M, heat wave 1 morning; HW1N, heat wave 1 afternoon; HW2N, heat wave 2 afternoon; HW3N, heat wave 3 afternoon; R1N, day 1 recovery from heat stress afternoon; CR2N, control plants parallel to the R2N time point afternoon; R2N, day 2 recovery from heat stress afternoon. Blue shading, control conditions; Pink shading, heat conditions. *A.t*, *Arabidopsis thaliana*; *A.h*, *Anastatica hierochuntica*; TPM, transcripts per kilobase million.

#### Validation of "between species" RNA-seq analyses

The above comparisons of gene expression between the two species utilized DeSeq2 (Love et al., 2014) as a normalized measure of gene expression and to identify differentially expressed genes. DeSeq2 normalizes read counts for different sequencing depths between samples. However, when dealing with two different species, several other factors can affect direct comparison of expression levels between orthologs including whether a few highly expressed genes constitute a large proportion of the sequenced transcripts, as well as differences in gene numbers and orthologous transcript length (Zhou et al., 2019; Zhao et al., 2020). Therefore, we performed several further analyses to validate our results. Figure S4A shows that there was no significant difference between the species in the proportion of the top 10 most highly expressed genes out of the total transcripts sequenced across all treatments (*A. thaliana*, ~7% to 12%, and *A. hierochuntica*, ~10% to 15% of the total sequenced transcripts). We further re-normalized our raw read count data (normalized for transcript length) using a new between-species method that applies Scale-Based Normalization (SCBN) to the most conserved orthologs, thereby obtaining a scaling factor that minimizes the false discovery rate of differentially expressed genes (Zhou et al., 2019). Applying SCBN to the 109 most conserved orthologs between *A. thaliana* and *A. hierochuntica* (Dataset S23) and using the scaling factor to correct normalized gene counts, we obtained similar comparative basal expression results as observed with DeSeq2 (Fig. S4B). Finally, QPCR analysis confirmed the RNA-seq fold-change gene expression patterns of four selected *A. thaliana* and *A. hierochuntica* genes (Fig. S4C). These genes included *AKS2,* a gene found to be positively selected in the "all extremophyte species" run (Table 1), two genes involved in abiotic stress responses (*ELIP1* and *HSP17.6II*; Sun et al., 2001; Rizza et al., 2011), and an *A. thaliana*-specific and an *A. hierochuntica*-specific Taxonomically Restricted Gene (*A. hierochuntica* ID: TRINITY_DN7044_c0_g2_i1; AGI: At2g07719; Methods S1). Additionally, we selected an ethylene signaling gene *CPR5* (Wang et al., 2017) that exhibited lower basal expression in *A. hierochuntica* than in *A. thaliana* in the RNA-seq analysis and confirmed this result via absolute QPCR (Fig. S4D).

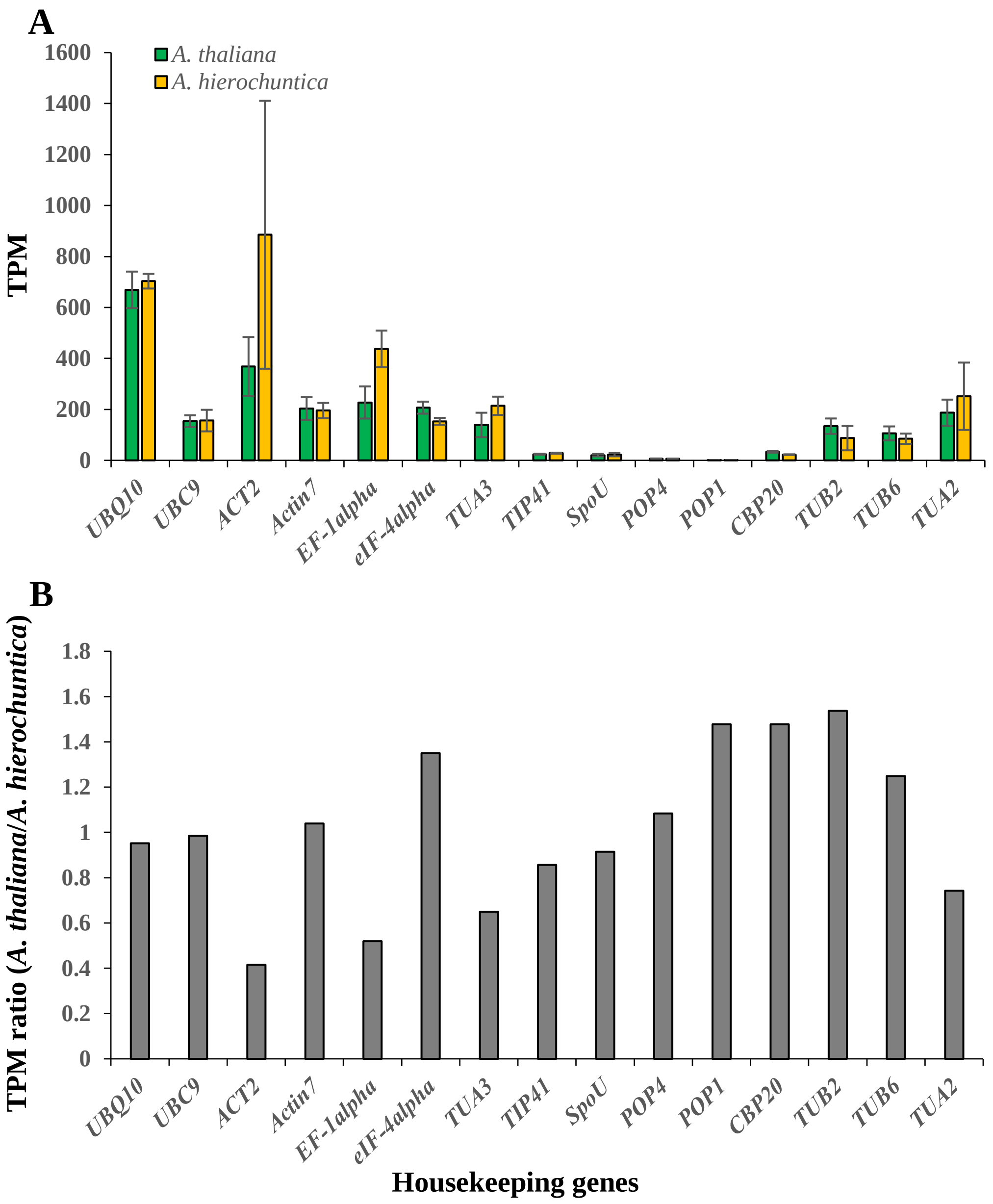


**Figure S5. Basal (control) expression of 15 orthologous *A. thaliana* and *A. hierochuntica* housekeeping genes.** (A) RNA-seq-based expression (see Methods S1); (B) Ratio of *A. thaliana*:*A. hierochuntica* expression.

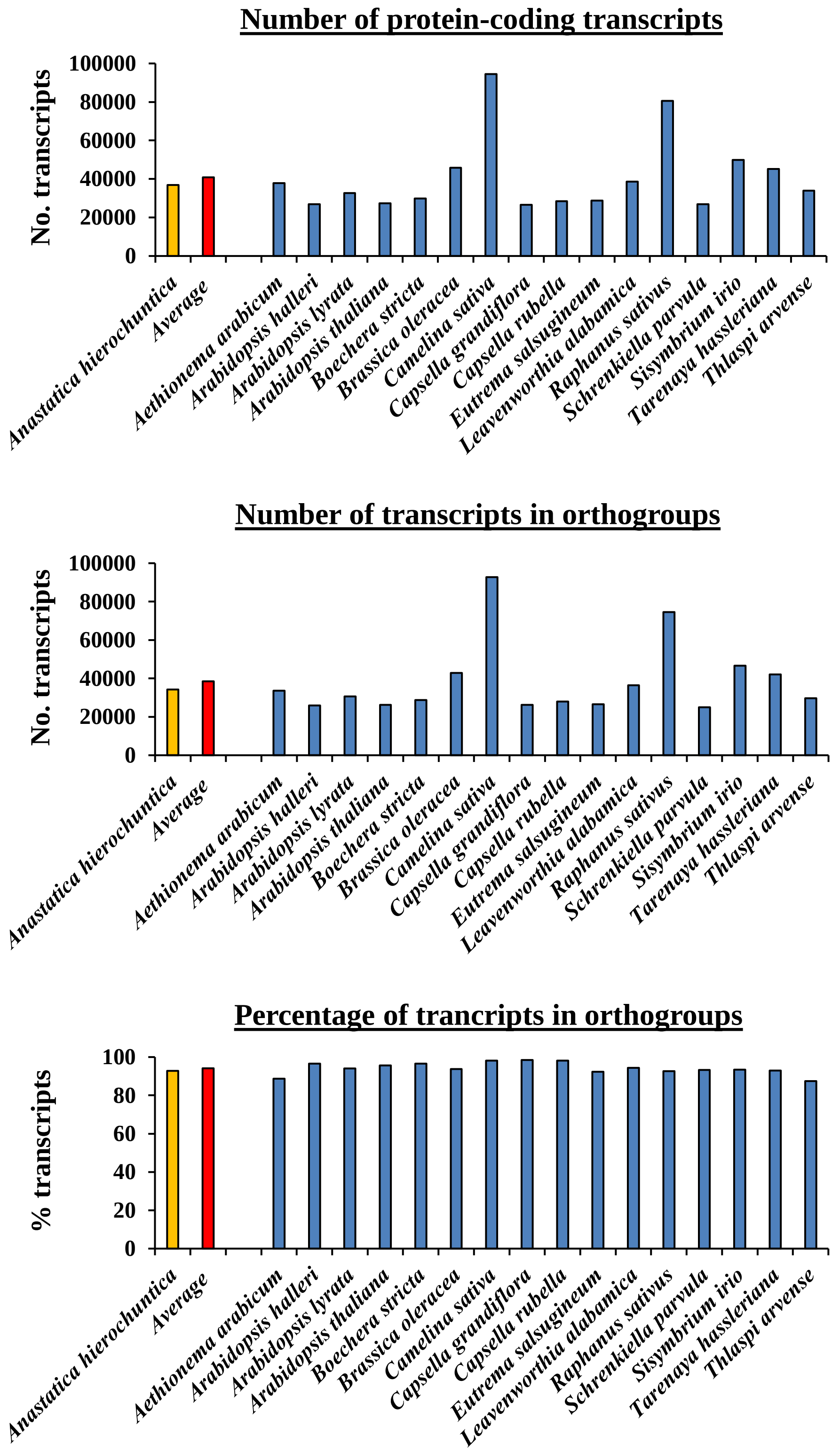


**Figure S6. Number of protein-coding transcripts and comparative ortholog group composition for species use to detect positively selected genes.** Data was generated using the OrthoFinder software (Emms and Kelly (2019), Genome Biol 20: 238).

### Biological Process

| | | | | |
|---|---|---|---|---|
| 11 | 9 | 8 | 5 | Anatomical structure development (GO:0048856) |
| 21 | 19 | 16 | 14 | Biological regulation (GO:0065007) |
| 29 | 29 | 21 | 22 | Biosynthetic process (GO:0009058) |
| 29 | 28 | 21 | 21 | Cellular biosynthetic process (GO:0044249) |
| 15 | 24 | 17 | 14 | Cellular macromolecule biosynthetic process (GO:0034645) |
| 24 | 34 | 24 | 24 | Cellular macromolecule metabolic process (GO:0044260) |
| 8 | 12 | 10 | 11 | Cellular protein metabolic process (GO:0044267) |
| 11 | 13 | 9 | 8 | Developmental process (GO:0032502) |
| 16 | 24 | 19 | 14 | Gene expression (GO:0010467) |
| 15 | 24 | 17 | 14 | Macromolecule biosynthetic process (GO:0009059) |
| 25 | 36 | 26 | 28 | Macromolecule metabolic process (GO:0043170) |
| 10 | 13 | 8 | 8 | Multicellular organismal development (GO:0007275) |
| 10 | 13 | 8 | 8 | Multicellular organismal process (GO:0032501) |
| 24 | 28 | 23 | 19 | Nitrogen compound metabolic process (GO:0006807) |
| 19 | 23 | 19 | 15 | Nucleobase, nucleoside, nucleotide and nucleic acid metabolic process (GO:0006139) |
| 9 | 12 | 12 | 15 | Protein metabolic process (GO:0019538) |
| 18 | 19 | 12 | 12 | Regulation of biological process (GO:0050789) |
| 11 | 14 | 9 | 8 | Regulation of biosynthetic process (GO:0009889) |
| 11 | 14 | 9 | 8 | Regulation of cellular biosynthetic process (GO:0031326) |
| 13 | 14 | 9 | 10 | Regulation of cellular metabolic process (GO:0031323) |
| 18 | 18 | 11 | 12 | Regulation of cellular process (GO:0050794) |
| 11 | 15 | 9 | 8 | Regulation of gene expression (GO:0010468) |
| 11 | 14 | 9 | 8 | Regulation of macromolecule biosynthetic process (GO:0010556) |
| 12 | 15 | 9 | 8 | Regulation of macromolecule metabolic process (GO:0060255) |
| 13 | 15 | 9 | 10 | Regulation of metabolic process (GO:0019222) |
| 11 | 13 | 9 | 8 | Regulation of nitrogen compound metabolic process (GO:0051171) |
| 11 | 13 | 9 | 8 | Regulation of nucleobase, nucleoside, nucleotide and nucleic acid metabolic process (GO:0019219) |
| 12 | 14 | 9 | 8 | Regulation of primary metabolic process (GO:0080090) |
| 11 | 13 | 9 | 8 | Regulation of transcription (GO:0045449) |
| 8 | 14 | 8 | 6 | Response to chemical stimulus (GO:0042221) |
| 6 | 10 | 5 | 5 | Response to organic substance (GO:0010033) |
| 14 | 20 | 14 | 11 | Response to stimulus (GO:0050896) |
| 9 | 11 | 12 | 6 | RNA metabolic process (GO:0016070) |
| 11 | 16 | 11 | 9 | Transcription (GO:0006350) |
| 7 | 8 | 6 | | Post–embryonic development (GO:0009791) |
| 9 | 10 | 8 | | Response to stress (GO:0006950) |
| 5 | | | | Anatomical structure morphogenesis (GO:0009653) |
| 5 | | | | Monocarboxylic acid metabolic process (GO:0032787) |
| | 5 | | | Macromolecule modification (GO:0043412) |
| | 5 | | | Protein modification process (GO:0006464) |
| | | 5 | | Lipid biosynthetic process (GO:0008610) |
| 7 | 5 | | | Cellular component biogenesis (GO:0044085) |
| 8 | 7 | | | Establishment of localization (GO:0051234) |
| 8 | 7 | | | Transport (GO:0006810) |
| 6 | | | 6 | Cellular nitrogen compound metabolic process (GO:0034641) |
| 7 | | | 6 | Heterocycle metabolic process (GO:0046483) |
| 8 | | | 7 | Localization (GO:0051179) |
| | 5 | 5 | | Translation (GO:0006412) |
| | | | | Carboxylic acid biosynthetic process (GO:0046394) |
| | | | | Cell growth (GO:0016049) |
| | | | | Cellular component morphogenesis (GO:0032989) |
| | | | | Cellular developmental process (GO:0048869) |
| | | | | Cellular response to stimulus (GO:0051716) |
| | | | | Cellular response to stress (GO:0033554) |
| | | | | Embryonic development (GO:0009790) |
| | | | | Embryonic development ending in seed dormancy (GO:0009793) |
| | | | | Fruit development (GO:0010154) |
| | | | | Growth (GO:0040007) |
| | | | | Negative regulation of biological process (GO:0048519) |
| | | | | Organic acid biosynthetic process (GO:0016053) |
| | | | | Regulation of anatomical structure size (GO:0090066) |
| | | | | Regulation of cell size (GO:0008361) |
| | | | | Regulation of cellular component size (GO:0032535) |
| | | | | RNA biosynthetic process (GO:0032774) |
| | | | | Seed development (GO:0048316) |
| | | | | Transcription, DNA–dependent (GO:0006351) |
| 5 | | | | Cellular nitrogen compound biosynthetic process (GO:0044271) |
| 5 | | | | Cofactor biosynthetic process (GO:0051188) |
| 5 | | | | Cofactor metabolic process (GO:0051186) |
| 5 | | | | Intracellular transport (GO:0046907) |
| | 5 | | | Post–translational protein modification (GO:0043687) |
| | 7 | | | Response to abiotic stimulus (GO:0009628) |
| | 5 | | | Response to abscisic acid stimulus (GO:0009737) |

40
30
20
10
0

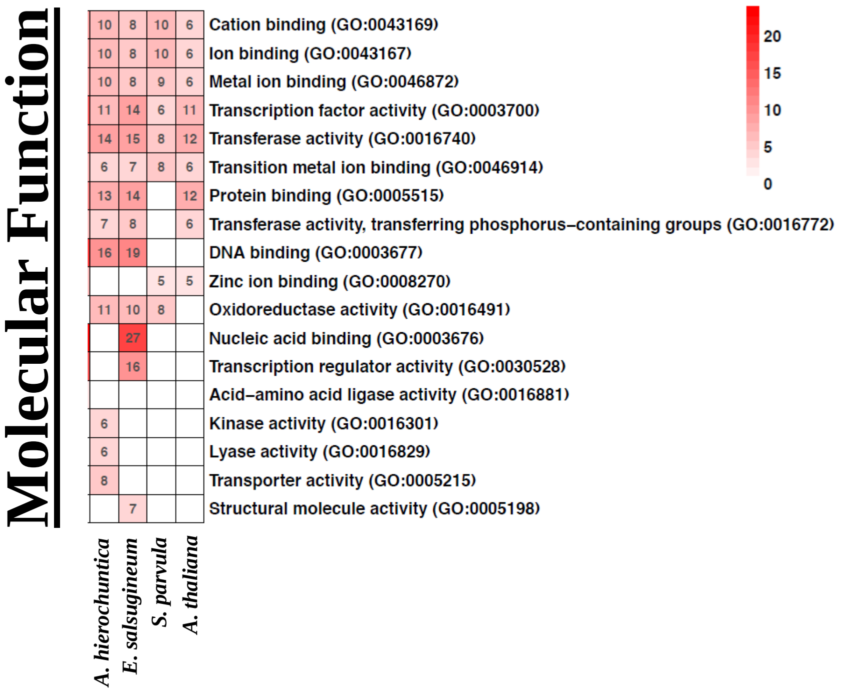


**Figure S7. GO-term enrichment analysis of positively selected genes.** Significantly (q-value < 0.05) enriched GO terms (biological process and molecular function) were compared among the five positive selection analyses and are represented in a heatmap. GO terms were assigned based on the *A. thaliana* ortholog sequence, or closest paralog. The *A. thaliana* genome served as the background for the enrichment analysis which was performed using the AgriGO online server (http://bioinfo.cau.edu.cn/agriGO/analysis.php). The red color intensity corresponds to the number of positively selected genes assigned with that GO term (the numbers are indicated within the cells). Cells with a white color correspond to GO terms that were not significantly enriched. For visualization and interpretation purposes, GO terms with > 2000 genes in the *A. thaliana* genome were excluded. Additionally, GO terms were clustered together if they share >50% of their gene set, and the GO term with the lowest q-value per cluster was included in the heatmap (Full enriched GO term lists can be found in Supplemental Tables S8-S12).

.
