## Supplemental Methods S1 for "Positive Selection and Heat-Response Transcriptomes Reveal Adaptive Features of the Brassicaceae Desert Model, *Anastatica hierochuntica*"

### Plant material and growth conditions

$F_4$ generation *A. hierochuntica* seeds descended from a single seed from plants originally collected in the Negev Desert (Nahal Hayun, 30.191424N and 35.009926E), Israel, were used in this study.

Seeds for plants used to extract RNA for sequencing of the *de novo* reference transcriptome were germinated and grown on nutrient agar plates for 5 d in the growth room (16 h light (150 µmol photons $m^{-2}$ $s^{-1}$)/8 h dark; 22 °C), as described in Eshel et al. (2017). Plant material was prepared for sequencing on two platforms: (i) For Illumina sequencing, plate-grown seedlings were harvested and snap-frozen in liquid nitrogen; (ii) For Roche 454 sequencing, plate-grown 5 day-old seedlings were transferred to pots containing autoclaved *A. thaliana* soil growth medium (Weizmann Institute of Science), irrigated to field capacity with 1 g $l^{-1}$ 20-20-20 NPK + micronutrients solution (Haifa Chemicals), and kept in the growth room until plants developed four true fully-expanded leaves. These plants were then treated with the following conditions: (a) Control (field-capacity, 22 °C); (b) Drought stress (25% field capacity for 1 week); (c) Salt shock (200 mM NaCl in the fertilizer solution), harvested after 1, 3 and 6 h; (d) Heat shock (45 °C), harvested after 0.5, 1, and 2 h. Roots, shoots and flowers (where available) from these soil-grown plants, were harvested separately and snap-frozen in liquid nitrogen. In addition to these soil-grown samples, mature seeds, from the same $F_4$ generation seed stock, were imbibed in $H_2O$ for 8.5 h, and then snap-frozen in liquid nitrogen.

For RNA-seq heat stress experiments, *A. thaliana* and *A. hierochuntica* were germinated and grown on nutrient agar plates according to Eshel et al. (2017). Seedlings were grown on plates until cotyledons were fully expanded before transfer to 7 cm x 7 cm x 8 cm pots containing *Arabidopsis* nitrogen-less soil (Weizmann Institute of Science; 70% fine peat [1–10mm], 30% perlite 4) irrigated to field capacity with a custom-made fertilizer solution (5 mM $KNO_3$, 2 mM $MgSO_4$, 1 mM $CaCl_2$ x $2H_2O$, 10 mM $KH_2PO_4$ [pH 6.0, adjusted with KOH] plus MS micronutrients (Murashige and Skoog, 1962). Flats containing pots were placed in the growth room under the same conditions as for the plate experiments. Flats were covered with plastic domes for 1 to 2 d, which were then gradually removed to allow seedlings to harden. Each day, pots were shuffled so that all plants received equal illumination and to remove shelf position effects. Plants were irrigated alternatively every 3 d with either fertilizer solution or water in order to maintain constant nutrient concentrations. After 6 d in the growth room, uniform plants were transferred to two growth chambers (KBWF 720, BINDER GmbH, Tuttlingen, Germany) (16 h light/8 h dark; 23 °C; 60% relative humidity) for heat treatments. The light/dark transitions at the beginning and end of the day comprised 0.5 h at 100 µmol photons $m^{-2}$ $s^{-1}$ and 0.5 h at 150 µmol photons $m^{-2}$ $s^{-1}$ to mimic sunrise and sunset. Light intensity for the remaining 14 h was 250 µmol $m^{-2}$ $s^{-1}$. Plants were allowed to acclimate for 4 d and were moved randomly every day between the chambers, to avoid chamber effects. At day 10 after transfer to soil, (*A. hierochuntica* plants had two true leaves and *A. thaliana* had six true leaves), heat treatment was initiated in one chamber, keeping the other chamber as the control (23 °C). The heat treatment included 3 d at 40/25 °C, day/night temperatures, followed by two days of recovery at control conditions (Fig. 2A). Similar to the 1 h light transition, the temperature was also gradually increased/decreased for 1 h, between the light/dark states, to reach the appropriate temperatures. Plants from both chambers were harvested at eight time points (Fig. 2A), either in the morning (1.5 h after the onset of the light/heat period) or at midday (7 h after the onset of the light/heat period). For each condition, three biological replicates comprising 6 pooled plants per replicate (27 samples per species) were used for downstream analyses.

**Reference transcriptome sequencing, assembly and annotation**

To generate a high-quality *A. hierochuntica* reference transcriptome that maximizes coverage of genes contained in the genome, we sequenced RNA pooled from multiple plant organs (root, shoot, flower, seeds), at different developmental stages (early seedling stage, and mature plants before and after anthesis), and under control and stress conditions (heat, drought and salinity).

For Illumina sequencing, a cDNA library was prepared using the TruSeq Stranded mRNA Sample Preparation kit (Illumina, San Diego, CA), according to the manufacturer's instructions. The library was then sequenced on Illumina Genome Analyser IIx (GAIIx), where 36,666,369 single-end, quality filtered, unaligned 76 nucleotide (nt) long raw reads, were generated with CASAVA 1.8 software. Reads containing adapter sequences were discarded, resulting in 36,369,571 filtered reads.

For Roche "454" sequencing, a normalized cDNA library was constructed using the Evrogen SMART technology cDNA synthesis service (Evrogen, Moscow, Russia) to reduce abundant RNAs such as Rubisco, and therefore enable detection of rare RNAs. The library was sequenced with a full picotiter plate on a 454 GS-FLX sequencer with Titanium reagents (Roche Applied Science, Indianapolis, IN, USA), yielding 862,788 raw reads (average read length of 315 nt). Read quality was monitored using FastQC (Babraham Institute, http://www.bioinformatics.babraham.ac.uk/projects/fastqc/). Raw reads were trimmed and filtered using the CLC Genomic Workbench (CLC bio, Inc.) with the following parameters: (i) Quality limit = 0.05; (ii) Ambiguous limit = 2; (iii) Minimum number of nucleotides in reads = 30; (iv) Adapter removal. To remove potential contaminant sequences, human, mouse, mosquito, bacteria, archaea and viruses sequence databases were screened using DeconSeq (Schmieder and Edwards, 2011) with alignment coverage length and identity thresholds of 95% and 94%, respectively. The potential contaminants were further BLASTN against the RefSeq_rna, and the TAIR10 databases, and searched using MEGABLAST against the nr databases, where reads with significant homology (e-value < $1e^{-5}$) to a plant sequence were retained. To account for potential inherent homopolymer insertion/deletion sequencing errors in the "454" reads, Illumina reads were used for error correction using the Blue error-correction algorithm (Greenfield et al., 2014). Altogether 724,653 high quality "454" reads were obtained. All Illumina and 454 reads were deposited in the SRA database under NCBI BioProject PRJNA731383.

The reference transcriptome was assembled using a hybrid assembly approach that utilized both Illumina and 454 reads. Briefly, the Illumina reads were first assembled using the Trinity software package (Grabherr et al., 2011). The resulting Trinity contigs were then shred into 700 bp fragments ("pseudo-reads"), with a 200 bp overlap between adjacent fragments, using the fasta2frag.tcl script, from the MIRA4 package (Chevreux et al., 1999; Chevreux et al., 2004). Both the shredded contigs and the 454 reads were assembled together using the Newbler assembler v.2.6 (Margulies et al., 2005) to generate the *A. hierochuntica* reference transcriptome comprising 36,871 assembled high-confidence transcripts (Figure S1).

Transcriptome completeness was assessed using the Benchmarking Universal Single-Copy Orthologs (BUSCO) tool (Simao, et al., 2015). TransDecoder (Haas et al., 2013) was used for predicting coding sequences in the Newbler-assembled transcripts. In order to assess the utility of the reference transcriptome for gene expression quantification, the Illumina reads were mapped to the reference *A. hierochuntica* transcriptome using the Bowtie read aligner program (Langmead et al., 2009), with the seed length set to 40 nt, and the maximum mismatches allowed within the seed length set to one.

For functional annotation, transcripts were searched for homolog sequences in other species using BLAST (e-value ≤ 1e$^{-5}$ cutoff) against the *A. thaliana* (TAIR10) coding sequences (CDS) database, a custom-made *Brassicaceae* CDS database (for included species see Dataset S12), and the curated, non-redundant RefSeq_protein and RefSeq_rna databases. Transcripts were also BLAST against the ncRNA databases, CANTATAdb (Szcześniak et al., 2015) and NONCODE2016 (Zhao et al., 2015). For these transcripts, ncRNA structures were also predicted using the online tool RNAcon (Panwar et al., 2014; http://crdd.osdd.net/raghava/rnacon/). Transcripts were also mapped to the Kyoto Encyclopedia of Genes and Genomes (KEGG) pathway database using the KAAS server (Moriya et al., 2007), with the bi-directional best-hit method against plant species (Zhang and Leong, 2010). The PlantTFcat server (Dai et al., 2013) was used for identifying regulatory genes (transcription factors (TFs), transcriptional regulators (TRs) and chromatin regulators (CRs). Transcripts were also searched for InterPro protein signatures using the InterProScan function (with the following applications: BlastProDom, FPrintScan, HMMPIR, HMMPfam, HMMSmart, HMMTigr, ProfileScan, HAMAP, PartterScan, SuperFamily, SignalPHMM, TMHMM, HMMPanther, Gene3D, Phobius and Coils) within the Blast2GO program (Conesa et al., 2005).

| **Functional annotation procedure** | **Total annotated transcripts** |
|---|---|
| Coding sequence prediction | 28,729 (78%) |
| InterPro signatures (InterProScan) | 27,511 (75%) |
| KEGG pathways (KAAS) | 9,010 (24%) |
| Plant regulatory gene families (PlantTFcat) | 4,043 (11%) |
| Best hit to *Arabidopsis thaliana* TAIR10 (BLASTN) | 25,386 (69%) |
| Best hit to *Brassicaceae* species (BLASTN) | 29,775 (81%) |
| Best hit to RefSeq_rna (BLASTN) | 33,263 (90%) |
| Best hit to RefSeq_protein (BLASTX) | 33,232 (90%) |
| Total annotated | 35,472 (96%) |

#### Identification of *A. thaliana* and *A. hierochuntica* taxonomically restricted genes (TRGs)

*A. thaliana* and *A. hierochuntica* transcriptomes were translated into ORFs and compiled into protein databases for the two species using TransDecoder (Haas et al., 2013). Reciprocal

BLASTp (ver. 2.10.1) was performed against the protein databases of 25 plant species with the following parameters: E-value ≤ $1e^{-5}$, Max. sequence target = 1. After compiling the transcripts that were not annotated, we performed BLASTn (ver 2.10.1) using the same parameters as above against the *A. thaliana* genome to further filter out transcripts which might have a putative function.

**RNA-seq of heat responsive *A. thaliana* and *A. hierochuntica* transcriptomes**

RNA-seq libraries were prepared with the Illumina 'TruSeq Stranded mRNA Sample Prep Kit (Illumina), multiplexed and pooled for each species separately (27 samples per species) and sequenced across 9 and 11 lanes for *A. thaliana* and *A. hierochuntica,* respectively. Sequencing was performed using an Illumina HiSeq2500 sequencer to generate 100nt single-end reads. In addition, a pooled sample of equal amounts of RNA from all *A. hierochuntica* samples, was prepared and sequenced generating 160nt paired-end reads. Fastq files were generated and demultiplexed with the bcl2fastq v2.17.1.14 Conversion Software (Illumina), while the FastQC program was used to monitor read quality. RNA-seq reads were deposited in the SRA database under NCBI BioProject PRJNA731383.

To estimate transcript abundance, 27 RNA-Seq *A. hierochuntica* fastq files were mapped to the reference transcriptome, while 27 RNA-Seq *A. thaliana* fastq files were mapped to the TAIR10 33,602 cDNA representative sequences. This was achieved using the Trinity align_and_estimate_abundance.pl script, which applies the RSEM program (Li and Dewey, 2011) with the Bowtie aligner. RSEM calculates the transcripts per kilobase million (TPM) normalized gene expression estimations, TPM values were used for individual gene expression graphs, while $\log_{10}$ transformed TPM values were used for the PCA analysis.

For statistical differential expression tests, count data of uniquely mapped reads were calculated per transcript from the Bowtie output using a custom python script, and were further analyzed using the R package DESeq2 (Love et al., 2014). In order to test for differences between species, a subset of 17,962 Bi-directional Best BLASTN hits (orthologs) between the two species, was used. DESeq2 normalizes the count data of each library by a size factor to account for different sequencing depths between the libraries. However, when comparing the expression levels between different species, it is essential to also normalize the read counts by the transcript length. Therefore, instead of inputting DESeq2 with the read counts per transcript, we first divided these values by the transcript length (in kilobases) and rounded the values to input DESeq2 with discrete numbers that fit a Poisson distribution.

One inherent problem in analyzing expression data (or any other type of multidimensional data) is that the variance increases with the mean. Therefore, the most highly expressed genes will dominate the differential expression analysis. One common solution is to use logarithmic transformation but this approach is prone to dominance of the lowest expressed genes, which will show the strongest differences between samples (see DESeq2 manual, http://www.bioconductor.org/help/workflows/rnaseqGene/#time-courseexperiments). Therefore, DESeq2 uses the regularized-logarithm transformation (rlog) to stabilize the variance in count data across the mean, thereby improving the dispersion estimation for both high and low expressed genes, and reducing the false discovery rate in calling differential expression (Love et al., 2014).

For analysis modes of expression and expression of genes associated with specific functions (Figs. 3A, 4 and 5B), genes were assigned GO terms for their respective categories using the BiNGO app of the Cytoscape software (Maere et al., 2005). The GO terms used for cell cycle were: ‘cell cycle’ (GO:0007049), ‘cytokinesis’ (GO:0000910), ‘regulation of cellular division’ (GO:0051302), ‘cell division’ (GO:0051301); The GO terms used for photosynthesis were: ‘photosynthesis’ (GO:0015979), ‘photorespiration’ (GO:0009853), ‘electron transport chain’, (GO:0022900) ‘light reaction’ (GO:0019684), ‘oxidation reduction’ (GO:0055114); The GO terms used for abiotic stress were: ‘response to water deprivation’ (GO:0019684), ‘resp. to heat’ (GO:0009408), ‘resp. to temperature stimulus’ (GO:0009266), ‘resp. to reactive oxygen species’ (GO:0000302), ‘resp. to hydrogen peroxide’ (GO:0042542), ‘resp. to high light intensity’ (GO:0009644), ‘resp. to oxidative stress’ (GO:0006979), ‘resp. to radiation’ (GO:0009314), ‘resp. to. abscisic acid stimulus’ (GO:0009737), ‘resp. to osmotic stress’ (GO:0006970), and ‘regulation of response to stress (GO:0080134)’.

For functional clustering of GO-terms in Fig. 5C, the GOMCL algorithm (Wang et al., 2020) was used to reduce redundancy of the Gene Ontology. GO terms that share the majority (>50%) of genes among them were clustered with an inflation value of 3. Enriched GO terms with more than 1,000 genes in the *A. thaliana* genome were excluded.

### Validation of RNA-seq data

*Scale-based normalization (SCBN)*

To validate the differential gene expression analysis performed with DeSeq2, we re-normalized our RNA-seq read count data (normalized to transcript length) using the Scale-Based Normalization (SCBN) method that aids in removing systematic variation between different species (Zhou et al. 2019), For the first step - pinpointing highly conserved genes - we utilized the 17,962 *A. thaliana* and *A. hierochuntica* orthologous genes identified using the Agalma phylogenomics pipeline (see below "Phylogenomics and positive selection analysis"). Applying BLASTn (ver, 2.10.1) software revealed 109 highly conserved orthologs with an E-value ≤ 1e-$^{100}$, query coverage of ≥ 98%, and identical matches of ≥ 99%. The SCBN R package (http://www.bioconductor.org/packages/devel/bioc/html/SCBN.html) was then used on the 109 highly conserved genes to obtain a scaling factor of 0.9223461, which it then applied to the 17,962 orthologs to call 12,808 common differentially expressed genes ($p \leq$ 1e-$^{05}$) between the two species. To generate an approximate corrected gene count, the scaling factor was applied to each individual gene and the corrected average and median basal (control) expression is depicted in Fig. 9B.

*Real-time QPCR analysis*

Total RNA was extracted from whole shoots with TRIzol (38% Phenol (w/v), 0.8 M guanidine thiocyanate, 0.4 M ammonium thiocyanate, 0.1 M sodium acetate pH 5, 5% glycerol (v/v)) according to Rio et al. (2010). To remove residual genomic DNA, 7 □g of total RNA was treated with RNase-free PerfeCta DNase I (Quanta Biosciences, Inc., Gaithersburg, MD, USA) according to the manufacturer's instructions, and cDNA was synthesized from 1 µg of total RNA using the qScript™ cDNA Synthesis Kit (Quanta Biosciences). For amplification of PCR products, primers were designed using the NCBI Primer-BLAST tool (Ye et al., 2012), and analyzed for any secondary structure with the IDT OligoAnalyzer™ Tool (Dataset S24). QPCR was performed with the ABI PRISM 7500 Sequence Detection System (Applied Biosystems). Each reaction contained 5 µl Applied Biosystems™ Power SYBR® Green PCR Master Mix (Thermo Fisher Scientific Inc., Waltham, MA, USA), 40 ng cDNA, and 300 nM of each gene-specific primer. The QPCR amplification protocol was: 95 °C for 60 s, 40 cycles of 95 °C for 5 s (denaturation) and 60 °C for 30 s (annealing/extension). Data were analyzed using the SDS 2.3 software (Applied Biosystems).

To check the specificity of annealing of the primers, dissociation kinetics was performed at the end of each PCR run. All reactions were performed in triplicates. Relative quantification of target genes was calculated using the $2^{-\Delta\Delta C}$T method (Livak and Schmittgen, 2001), using *A. thaliana eIF4A1* and the *A. hierochuntica eIF4A1* ortholog as internal references. To ensure the validity of the $2^{-\Delta\Delta C}$T method, standard curves of 2-fold serial dilutions of cDNA were created and amplification of efficiencies of target and reference gene were shown to be approximately equal. For absolute quantification of basal expression, QPCR products were gel-purified (Gel/PCR DNA Fragments Extraction Kit [Geneaid Biotech Ltd, New Tapei City, Taiwan]), and quantified with a Nanodrop spectrophotometer. Fivefold serial dilutions of each PCR product were used to create a standard curve for determination of transcript copy number. As a loading control, the absolute transcript copy number of *eIF4A1*was also calculated and normalized to the highest *eIF4A1*level, which was assigned a value of 1. The target gene transcript copy number was then adjusted for loading differences by dividing by the normalized *eIF4A1* level.

**Clustering by Weighted Gene Correlation Network Analysis (WGCNA) and GO enrichment**

WGCNA was performed according to Langfelder and Horvath (2008) using R version 3.6.1 in RStudio (version 1.2.1335 (RStudio, Inc.)). WGCNA was set to using a weighted network analysis with Pearson correlations. The soft thresholding power (argument 'power' in function 'Adjacency') was set to 24 for *A. thaliana* and 30 for *A. hierochuntica*. Minimum module size (argument 'minClusterSize' in function 'dynamicMods') was set to 50 for both species, and module merging height cut (argument 'cutHeight' in function 'mergeCloseModules') was set to 0.30 for both species. Module expression profile heatmaps were produced from data created by WGCNA using the R package pheatmap (Kolde, 2012). A custom script was written to scale the expression data of each module's constituent genes relative to the other genes in the module so that expression pattern rather than expression level was visualized.

Gene Ontology (GO) enrichment analysis of the heat shock modules was performed using the Cytoscape (version 3.7.1) software app BiNGO (Shannon et al., 2003; Maere et al., 2005). For *A. thaliana*, a significance level of 0.05 was used, with an overrepresentation of ≥ 1.5. For *A. hierochuntica*, the respective *A. thaliana* orthologs were used. To correct for any high GO-term enrichment bias in *A. hierochuntica,* orthologs of the whole annotated *A. hierochuntica* transcriptome were tested for GO enrichment. The highest level of GO term enrichment was 1.38.

Since for *A. thaliana* GO terms are usually only considered enriched at ≥ 1.5-fold ($q \leq 0.05$), we corrected for any bias in the *A. hierochuntica* transcriptome by using a cutoff of ≥ 2.0 (1.5 * 1.38 ≈ 2.0)) with $q \leq 0.05$.

**Phylogenomics and positive selection analysis**

To identify positively selected genes, which are unique to *A. hierochuntica* or common to extremophyte Brassicaceae species, we used coding sequences of the *A. hierochuntica* reference transcriptome and 16 sequenced Brassicaceae species (Dataset S12) into the automated Agalma phylogenomics pipeline (Dunn et al., 2013).

The Agalma pipeline identified orthologous genes among these species by: (i) Identifying homologous genes among all input sequences from all the species using an all-by-all TBLASTX search followed by a Markov Clustering Algorithm (MCL) tool (Enright et al., 2002); (ii) For each homolog group, a peptide multiple sequence alignment (MSA) is produced using MAFFT with the E-INS-i algorithm (Katoh et al., 2005); (iii) The MSA is further used to build a maximum likelihood (ML) phylogenetic tree with RAxML v8.2.3 (Stamatakis, 2014). (iv) The homolog group tree is further pruned into maximally inclusive subtrees to define ortholog groups. MSAs of 13,806 ortholog groups with a sequence representation in at least 4 taxa were concatenated into a supermatrix for ML species tree search, using RAxML (with the PROTGAMMAWAG model of evolution, and 100 rapid bootstrap searches) under the WAG rate matrix (Whelan and Goldman, 2001), with gamma-distributed among-site rate variation.

To detect positive selection in the five extremophyte species, ortholog groups with sequence representation in at least 2 extremophytes, were selected to ensure sufficient statistical power (Anisimova et al., 2001). For each ortholog group, the peptide MSA was converted into the corresponding codon alignment using the pal2nal.pl program (Suyama et al., 2006), and the ML species tree was pruned using PHAST tree_doctor (Hubisz et al., 2011), to keep only sequence-represented taxa. Codon alignments together with pruned trees were further analyzed with the PAML v4.8, CODEML program (Yang, 1997; Yang, 2007), using the Branch-Site model. To test for positive selection, the tested branch(s) were labeled (foreground), and the log likelihood of two models (M1a and M2a), were calculated for each ortholog group. The difference between the two models is that in the M1a (null) model, the non-synonymous to synonymous rate ratio (dN/dS) is fixed to 1 (fix_omega = 1 and omega = 1), indicative of neutral selection, while in the M2a (alternative) model, the initial dN/dS ratio is set to 1, and is further estimated by the model (fix_omega = 0 and omega = 1). A Likelihood Ratio Test was performed (with $X^2$ distribution), to identify genes with log likelihood values significantly different between the two models, indicative of deviation from neutral selection. Ortholog groups with portion of sites in the foreground branches, that had an estimated dN/dS ratio greater than 1, were considered under positive selection. To account for multiplicity, a Benjamini–Yekutieli false discovery rate (FDR) correction (Benjamini and Yekutieli, 2001) was applied using the "qvalue" R package, with a $q$-value < 0.05 cutoff for a gene to be considered as positively selected. Sites under positive selection were identified using the empirical Bayes approach with a posterior probability $p > 0.95$.

The above procedure was repeated to identify positive selected genes in *A. hierochuntica*, and in the other extremophyte species. For each analysis, different branches on the tree were tested (labeled as foreground) compared with all other branches (background): (i) labeling the external branches of all five extremophyte species as the foreground (4,723 ortholog groups); (ii) labeling the *A. hierochuntica* external branch as the foreground (3,093 ortholog groups); (iii) labeling the *E. salsugineum* external branch as the foreground (4,457 ortholog groups); (iv) labeling the *S. parvula* external branch as the foreground (4,369 ortholog groups); and (v) labeling the *A. thaliana* external branch as the foreground (5,513 ortholog groups). *A. thaliana* was considered as a control/comparator species sensitive to abiotic stresses (Kazachkova et al., 2018). The Venn diagram comparing positive selected genes (Fig. 6B) was generated using an online tool: http://bioinformatics.psb.ugent.be/webtools/Venn/).

To further assess the functionality of the positively selected genes, Gene Ontology (GO) terms were assigned to each ortholog group based on *A. thaliana* GO annotation. In cases where an ortholog group did not contain an *A. thaliana* ortholog, the closest *A. thaliana* homolog (best BLASTP hit) was used. Significant positively selected genes were further tested for enriched GO terms (Fisher's exact test, with a $q$-value < 0.05 cutoff) using the online AgriGO tool (Du et al., 2010; http://bioinfo.cau.edu.cn/agriGO/analysis.php), where the *A. thaliana* genome served as the background. Enriched GO terms with more than 2,000 genes in the *A. thaliana* genome were excluded, as these are broad and less informative terms.
